# N-Cadherin Provides a Cis and Trans Ligand for Astrotactin that Functions in Glial-Guided Neuronal Migration

**DOI:** 10.1101/357541

**Authors:** Zachi Horn, Hourinaz Behesti, Mary E. Hatten

## Abstract

Prior studies demonstrate that Astrotactin (ASTN1) provides a neuronal receptor for glial-guided CNS migration. Here we report that ASTN1 binds N-cadherin (CDH2) and that the ASTN1:CDH2 interaction supports cell-cell adhesion. To test the function of ASTN1:CDH2 binding in glial-guided neuronal migration, we generated a conditional loss of *Cdh2* in cerebellar granule cells and in glia. Granule cell migration was slowed in cerebellar slice cultures after a conditional loss of neuronal *Cdh2*, and more severe migration defects occurred after a conditional loss of glial *Cdh2*. Expression of a mutant form of ASTN1 that does not bind CDH2, in granule cells, also slowed migration. Moreover, *in vitro* chimeras of granule cells and glia showed impaired neuron-glia attachment in the absence of glial, but not neuronal, *Cdh2*. Thus, *cis* and *trans* bindings of ASTN1 to neuronal and glial CDH2 form an asymmetric neuron-glial bridge complex that promotes glial-guided neuronal migration.

## INTRODUCTION

In cortical regions of mammalian brain, glial-guided neuronal migration directs postmitotic cells into neuronal layers, a process that underlies the formation of the cortical circuitry (1–3). The cerebellar cortex has long provided a key model for understanding the molecular basis of glial-guided migration, as granule cell precursors (GCPs) migrate from the external germinal layer (EGL) along the radial processes of Bergmann glia (BG) to a position deep to the Purkinje neuron, the sole output neuron of the cerebellar cortex (4). Correlated video and electron microscopy (EM) imaging of GCP migration along BG demonstrates that migrating neurons form a *puncta adherens* migration junction beneath the cell soma and extend a motile leading process in the direction of forward movement (5, 6). During migration, the neuron forms and releases the migration junction by a process that involves endocytosis of the receptor Astrotactin (ASTN1), which is expressed in neurons but not in glia (7). Molecular experiments demonstrate that the conserved polarity complex mPar6 regulates the cadence of locomotion by controlling the forward movement of the centrosome (8), as well as microtubule dynamics and acto-myosin motor function in the proximal aspect of the leading process (9), with the Rho GTPase Cdc42 controlling actin dynamics required for the polarity of the migrating GCP and for the formation of the migration junction with the glial fiber (10). While biochemical and genetic experiments have confirmed the key role of the neuronal guidance receptor ASTN1 in the migration junction (11–13), evidence is lacking on the glial ligand for ASTN1.

Cadherins are cell surface proteins composed of an adhesive extracellular domain and a cytoplasmic tail that links to the actin cytoskeleton through a complex of catenins. The extracellular domain allows cadherins to form lateral (*cis*) homodimers or mediate cell adhesion through *trans* homodimers. A large body of evidence demonstrates a key role for homophilic *trans* cadherin interactions in the formation and maintenance of *puncta adherens* junctions in the developing heart and neural tube (14), and in synapse formation (15, 16). In addition, disruption of the neural cadherin, N-cadherin (CDH2), leads to defects in neuronal migration during development of the cerebral cortex (17–22). Here we show that an asymmetric *cis* and *trans* complex of ASTN1 and CDH2 functions in neuronal migration. Conditional loss of glial CDH2 in mice impaired GCP migration *in vivo* and *ex vivo*, and perturbed the formation of a migration junction between GCPs and BG in cell-based assays. Moreover, CDH2-deficient GCPs expressing an ASTN1 variant that lacks the binding domain for CDH2 failed to migrate on CDH2-expressing glia. This suggests that ASTN1 in neurons, and CDH2 in neurons and glial fibers, form an asymmetric bridge complex that is required for glial-guided migration, and, more generally, that CDH2 might function as a heterophilic binding partner in the formation of other cell-cell junctions.

## RESULTS

### CDH2 is Expressed in the Migration Junction and Interacts with ASTN1

To investigate whether CDH2 interacts with ASTN1, we performed immunoprecipitation (IP) on protein lysates from postnatal day 7 (P7) mouse cerebella using an ASTN1 antibody. In this assay, we found that ASTN1 interacts with CDH2 (Fig. 1 A). Similarly, in HEK 293T cells, we detected co-IP of CDH2 with full length ASTN1 (ASTN1-FL) but not with a mutant variant of ASTN1, lacking a large portion of the C-terminal ectodomain (ASTN1-ΔCTD) that included the membrane attack complex/perforin (MACPF), fibronectin type III (FNIII), and annexin-like (ANX-like) domains (Fig. 1 B). Flow cytometry showed that ASTN1-FL and ASTN1-ΔCTD localized to the cell surface at similar levels (58 *%* and 49 %, respectively; Fig. S1). Thus, the extracellular C-terminus of ASTN1 forms a *cis* interaction with the ectodomain of CDH2.

**Fig. 1.**
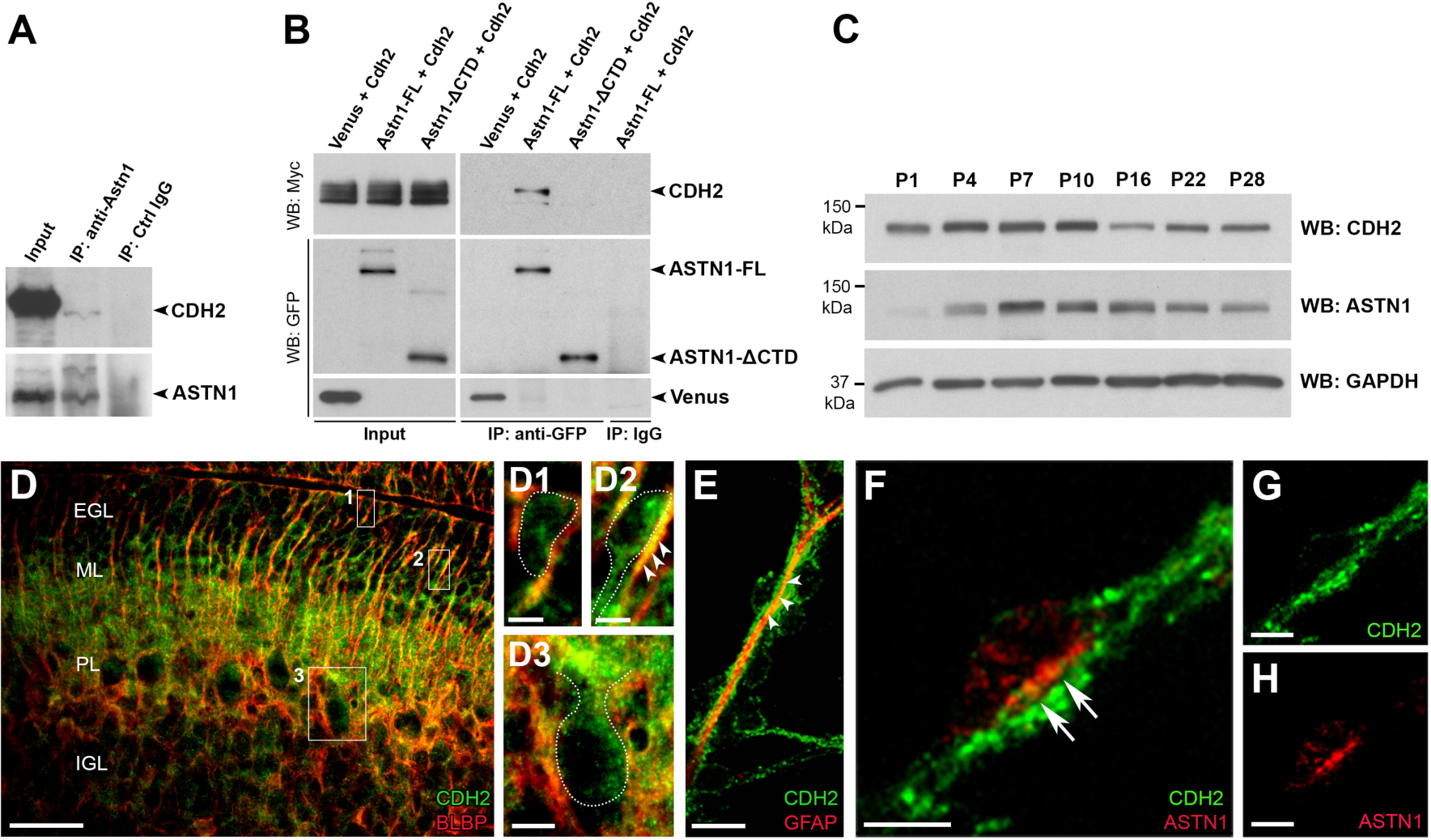
ASTN1 and CDH2 form *cis* interactions and co-localize in the migration junction. **(A)***In vivo* immunoprécipitation of ASTN1 in P7 whole cerebellar lysates, blotted with ASTN1 and CDH2 antibodies. ASTN1 was part of a protein complex with CDH2. **(B)** Western blots showing co-immunoprecipitation of ASTN1-Venus and CDH2-Myc in HEK 293T cells. CDH2 interacted with ASTN1-FL, but not with ASTN1-ΔCTD. **(C)** Developmental protein expression of CDH2 and ASTN1 in the cerebellum of postnatal mice (P1 - P28) by Western blot. CDH2 expression was highest between P4 - P10, decreased by P16 and reached a steady level at P22 - P28. ASTN1 expression increased after P1 and was highest at P7 - P10. Protein expression was compared to GAPDH levels. **(D, E)** Endogenous protein expression of CDH2 at P7 in sagittal mouse cerebellar sections **(D)** and in GCP/BG *in vitro* co-cultures **(E)**. CDH2 was expressed in GCPs in the EGL **(D1)**, migrating GCPs in the ML **(D2)** and in Purkinje cells **(D3)**, and co-localized with BLBP and GFAP in BG fibers (arrowheads in D2 and E). **(F - H)** GCP/BG co-cultures labeled with antibodies against CDH2 and ASTN1. CDH2 localized to neuronal processes, glial fibers and the migration junction beneath the neuronal soma. ASTN1 co-localized with CDH2 in the migration junction (arrows). EGL: external granule layer; ML: molecular layer; PL: Purkinje cell layer; IGL: internal granule layer. Scale bar represents 50 μm in (D), 5 μm in (D1, D2, F - H) and 10 μm in (D3, E).

We then used Western blotting (WB) and immunohistochemistry to examine the expression of CDH2 in migrating GCPs *in vivo*. WB of whole cerebellar lysates showed maximal levels of CDH2 in the early postnatal stages (P4 - P10), when ASTN1 expression is high (Fig. 1 C). In sections of early postnatal mouse cerebellum (P5 - P7), a CDH2 antibody labeled GCPs in the EGL, GCPs migrating across the molecular layer (ML), and mature granule cells (GCs) in the internal granule layer (IGL) as well as in the radial processes of BG stretching across the ML and in Purkinje cells (Fig. 1 D).

By immunostaining, CDH2 also localized to the migration junction, a *puncta adherens* junction between migrating GCPs (6), identified by their elongated profile and close apposition with the glial fiber (5, 23), and BG fibers in cultures of purified neurons and glia (Fig. 1 E - H). Antibodies against CDH2 intensely labeled the neuronal soma at the junction with the glial fiber and also stained the underlying glial fiber. In agreement with prior light and immuno-EM localization studies (11), antibodies against ASTN1 labeled the neuronal aspect of the migration junction (Fig. 1 F, H). Thus, ASTN1 and CDH2 co-localize to the migration junction of GCPs migrating along BG fibers.

### CDH2 and ASTN1 Form Heterophilic *Trans* Interactions

To analyze whether CDH2 interacts with ASTN1 in *trans* to promote cell adhesion, we used a classical S2 cell adhesion assay (24). For this assay, we transfected S2 cells with bicistronic expression constructs (25) of *Cdh2;GFP* or *Astn1;mCherry* cDNA and measured cell aggregation rates over two hours. Cells transfected with *Cdh2;GFP* formed aggregates within minutes, demonstrating a rapid homophilic *trans* binding of CDH2 (Fig. 2 A). In contrast, ASTN1-positive cells did not form homophilic aggregates over the two hour incubation period. ASTN1-positive cells did, however, form co-aggregates with CDH2-positive cells, indicating heterophilic *trans* binding between ASTN1 and CDH2 (Fig. 2 B). At two hours, CDH2-positive aggregates contained 11.7 ±1.2 % ASTN1-positive cells compared to 2.6 ±0.7 % mCherry-expressing control cells (*P* < 0.0001). Moreover, in contrast to the control cells, ASTN1-positive cells frequently integrated into the core of the aggregates. Taken together, these results confirmed earlier findings that ASTN1 does not promote cell adhesion through homophilic binding (26), and showed that CDH2 provides a *trans* ligand for ASTN1 that functions in cell-cell adhesion.

**Fig. 2.**
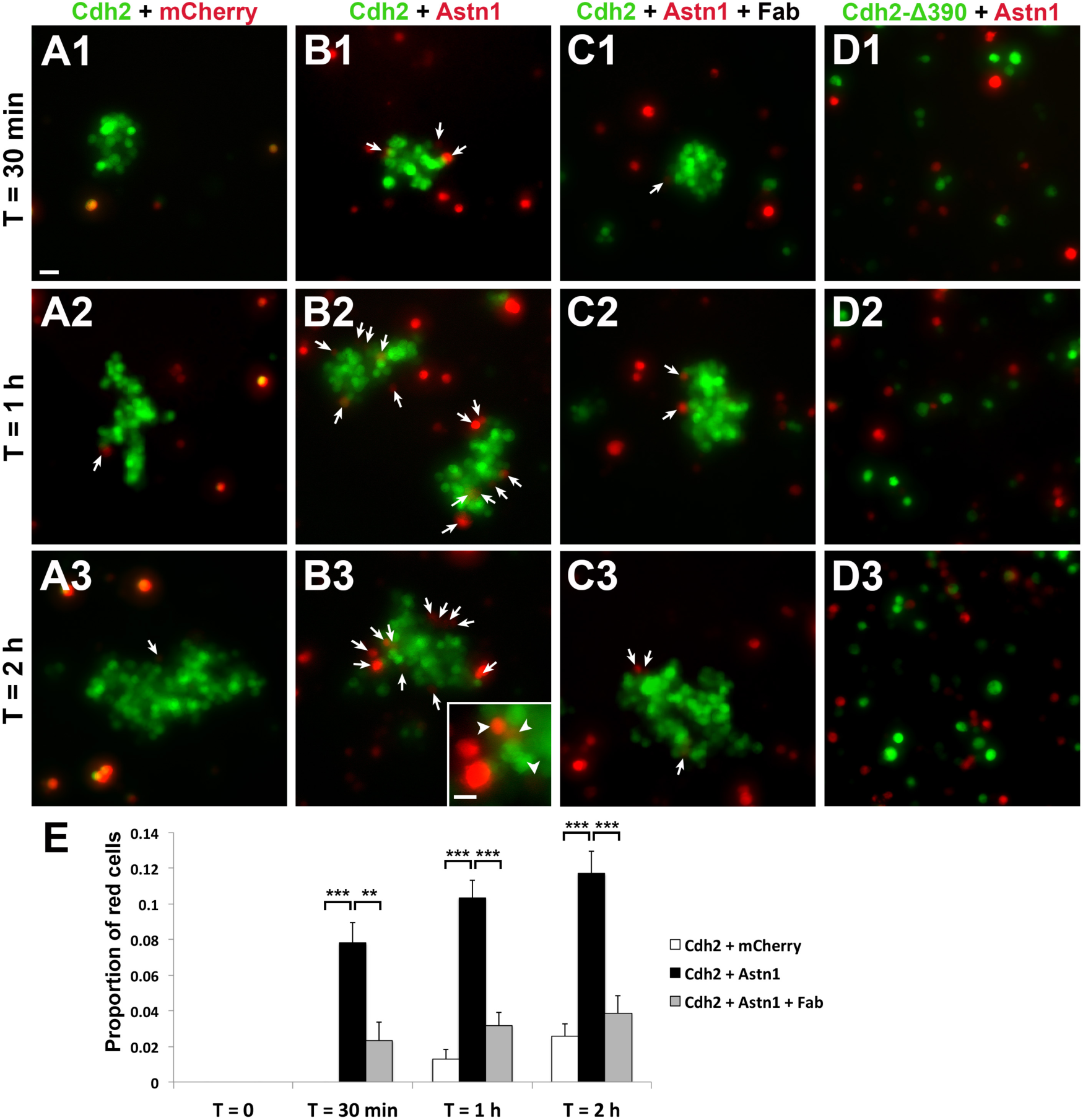
Heterophilic *trans* interactions of ASTN1 and CDH2. *Drosophila* S2 cell adhesion assays were prepared in four conditions: *Cdh2;GFP* + *mCherry;GFP* **(A)**, *Cdh2;GFP* + *Astn1;mCherry* **(B)**, *Cdh2;GFP* + *Astn1;mCherry* + ASTN1 Fab **(C)**, and *Cdh2-A390;GFP* + *Astn1;mCherry* **(D)**. ASTN1-positive cells were adhering to the CDH2-expressing aggregates after 30 min (arrows in **B1**), with more co-aggregation seen after 1 h (arrows in **B2**) and 2 h (arrows in **B3**), indicating heterophilic *trans* interactions. Significantly lower proportions of cells were adhering to the aggregates in the conditions with cells expressing control vector **(A)** or *Astn1;mCherry* blocked with ASTN1 Fab fragments **(C)**. The proportion of mCherry expressing cells in the CDH2;GFP-positive aggregates is quantified in **(E)**. Expression of CDH2∆390;GFP did not result in cell aggregation within 2 h **(D)**, demonstrating the importance of the cadherin ectodomain for homophilic and heterophilic interactions and cell adhesion. ** *P <* 0.01; *** *P* < 0.001. Scale bar represents 20 μm in (A - D) and 10 μm in inset in (B3).

To test the specificity of the heterophilic CDH2:ASTN1 *trans* interaction, we measured the aggregation of ASTN1 and CDH2-positive S2 cells in the presence of Fab fragments of an ASTN1 antibody raised against the C-terminus of ASTN1 (12). After addition of Fab fragments, heterophilic CDH2:ASTN1 *trans* cell adhesion was reduced to control levels (Fig. 2 C), suggesting that the C-terminus of ASTN1 is required for the interaction with CDH2. We then assessed the specificity of both homophilic CDH2:CDH2 and heterophilic CDH2:ASTN1 binding using a *Cdh2-Δ390* construct with a deletion of the extracellular domain of CDH2. In S2 cells expressing CDH2-Δ390, both homophilic adhesion and heterophilic adhesion with ASTN1-positive cells failed (Fig. 2 D), demonstrating a requirement for the ectodomain of CDH2 in *trans* cell adhesion.

### Cell-Specific Deletion of *Cdh2* in the Cerebellum

To provide a genetic model for the function of CDH2 in GCP migration in the developing mouse cerebellum, we generated a conditional knockout (cKO) of *Cdh2* by crossing a floxed *Cdh2* (*Cdh2*^*fl/fl*^) mouse line with a *NeuroD1-Cre* line to delete *Cdh2* in GCPs. In addition, we crossed the *Cdh2*^*fl/fl*^ mice with an *mGFAP-Cre* line to delete *Cdh2* in BG, or with or an *hGFAP-Cre* line to delete *Cdh2* in both GCPs and glia (27–29). Western blot analysis of lysates of GCPs and BG purified from each of the lines at P7 (30) confirmed the cell-specific deletion of *Cdh2* (Fig. S2 A). To examine the development of the cerebellum in each of these cKO lines, we first analyzed fixed sections of P7 cerebellum by Nissl staining (Fig. S2 B - E). Although the overall size of the cerebellum did not differ significantly in the three lines (*n* = 7 per genotype), defects in the foliation pattern of the cerebellum of *Cdh2*^*fl/fl*^ *;mGFAP-Cre* and *Cdh2*^*fl/fl*^*;hGFAP-Cre* mice were observed compared with controls. These defects included additional fissures in the ventral (I-III) lobes, with fewer fissures and irregularly shaped lobes in medio-dorsal (VI-VIII) areas.

### Loss of *Cdh2* in Granule Cells and/or Glia Has a Differential Effect on Migration

Immunostaining of cerebellar sections of the three cKO lines with NeuN, a marker for GCs, and with BLBP, a marker for BG, revealed striking differences in GCP migration and formation of the neuronal layers, especially in lines where BG or both BG and GCPs lacked *Cdh2* (Fig. 3). In all three lines, the density and organization of NeuN-positive GCPs in the EGL was identical to that seen in control mice. In *Cdh2*^*fl/fl*^*;NeuroD1-Cre* mice, the profile of GCPs migrating across the molecular layer was indistinguishable from controls and the overall laminar organization of the cerebellum appeared to be normal (Fig. 3 A, B). In contrast, NeuN-positive GCPs in the ML of both *Cdh2*^*fl/fl*^ *;mGFAP-Cre* and *Cdh2*^*fl/fl*^*;hGFAP-Cre* had rounded nuclei and many GCPs were located in the ML, suggesting slowed or stalled GCP migration (Fig. 3 C – F) Immunostaining for BLBP in these mice also revealed defects in positioning of BG cell bodies and radial patterning of BG fibers in some areas of the cerebellum (Fig. 3 D,F) Overall, the laminar patterning of the cerebellum of mice was disorganized, as the classic boundaries of the ML and IGL were uneven relative to controls. However, immunostaining with a Calbindin antibody showed no changes in the gross morphology or positioning of Purkinje cells (data not shown). Thus, a conditional loss of *Cdh2* in BG perturbed the migration of GCPs, the patterning of BG fibers and the formation of GC layers.

**Fig. 3.**
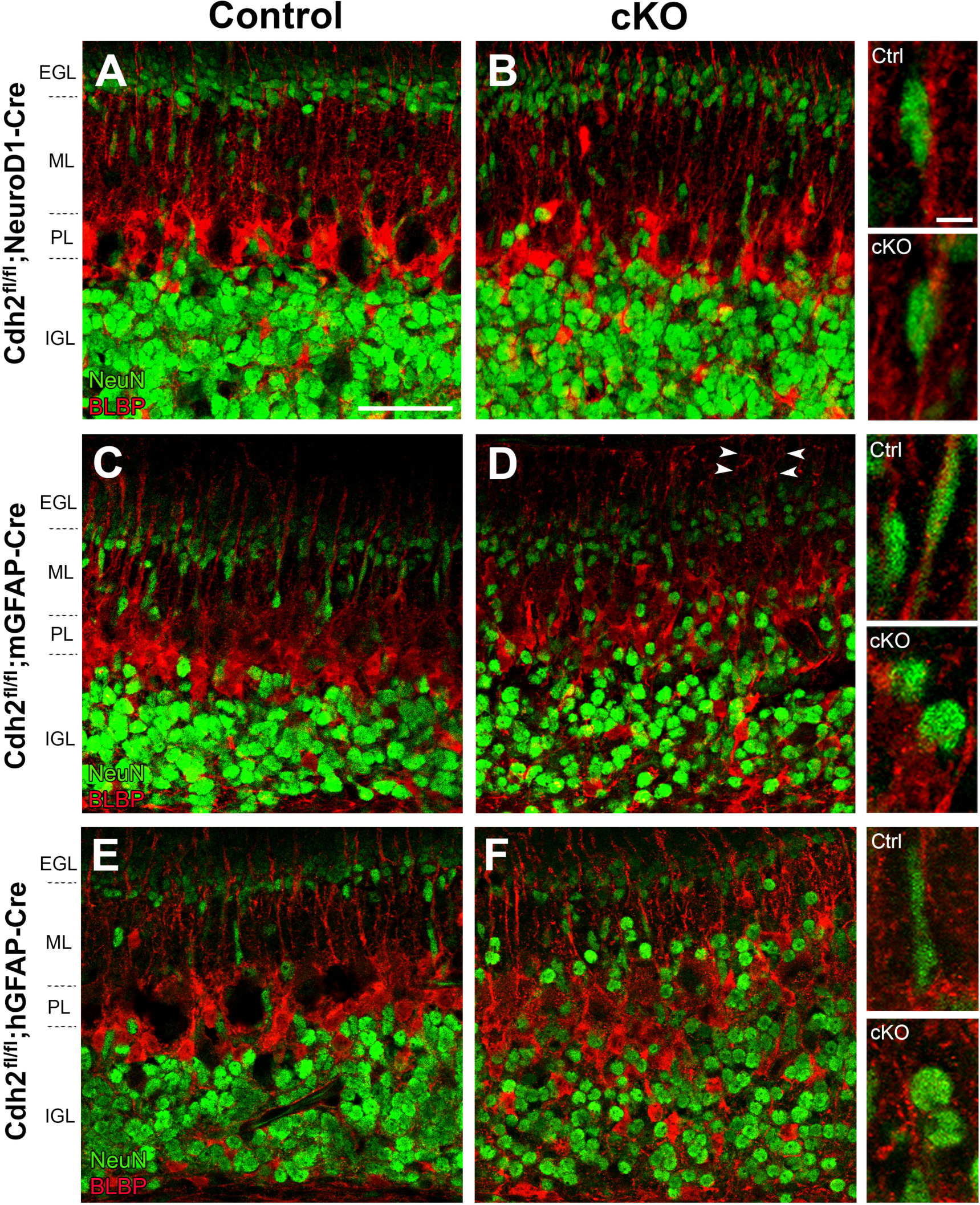
Neuronal migration in *Cdh2* cKO mice. Sagittal cerebellar sections of P7 *Cdh2*^*fl/fl*^ control mice **(A, C, E)** and *Cdh2* cKO littermates expressing *NeuroD1-Cre* **(B)**, *mGFAP-Cre* **(D)** or *hGFAP-Cre* **(F)**, and labeled with NeuN and BLBP antibodies. In control mice, NeuN-positive GCPs displayed elongated nuclei along BG fibers in the molecular layer (ML), indicating migrating cells. A similar phenotype is seen in mice with GCPs lacking *Cdh2* **(B)**. In contrast, a loss of *Cdh2* in BG **(D)** or in both GCPs and BG **(F)** resulted in GCPs with rounded nuclei and a stalled migration in the ML. In addition, abnormal radial patterning of BG fibers was observed in some areas (arrowheads in D). Insets show representative cells from each genotype. EGL: external granule layer; PL: Purkinje cell layer; IGL: internal granule layer. Scale bar represents 50 μm in (A - F) and 5 μm in insets.

To quantitate the rate of GCP migration along BG fibers, we performed BrdU birth-dating experiments, injecting BrdU at P5 and sacrificing the animals at P7 (*n* = 4 per genotype; Fig. S3). By BrdU labeling, the migration distance of GCs was reduced in *Cdh2*^*fl/fl*^*;NeuroD1-Cre* mice where 40 ±3.7 % of labeled GCs reached the IGL compared to 50 ±3.9 % in control (P = 0.035). However, GCP migration was dramatically reduced in the *Cdh2*^*fl/fl*^*;mGFAP-Cre* mice, where 33 ±4.4 % reached the IGL compared to 60 ±1.9 % in control (*P* = 0.040). Similarly, in the *Cdh2*^*fl/fl*^*;hGFAP-Cre* mice, 29 ±2.9 % of GCs reached the IGL compared to 55 ±4.4 % in control (*P* = 0.0002). The latter two lines also had a significantly higher proportion of BrdU-labeled cells in the ML (22 % and 21 % higher than control littermates, *P* = 0.02 and 0.0002, respectively). These findings suggest that the expression of CDH2 in BG fibers is required for GCP migration.

Since a decrease in the number of GCs reaching the IGL could also be due to changes in cell proliferation or cell death, we stained sections with antibodies to phosphohistone H3 and the apoptosis marker Caspase-3. We found no differences in proliferation or apoptosis in the three cKO lines (data not shown), suggesting that the lower proportion of cells in the IGL resulted from migration defects.

### Glial CDH2 is Essential for Granule Cell Migration in Organotypic Slice Cultures

To analyze the features of migrating GCPs in the three cKO lines in more detail, we used electroporation to express the fluorophore Venus in GCPs in P8 organotypic slices of cerebellar cortex and imaged labeled cells by spinning disc confocal microscopy (Fig. 4). In control slices, after 60 hours, Venus-positive GCs were observed in the inner EGL, ML and the outer portion of the IGL. Venus-positive cells in the inner EGL extended long parallel fiber axons, with labeled cells in the ML showing the bipolar morphology typical of migrating neurons with a leading process in the radial plane (Fig. 4 A, C, E). Although the polarity and overall morphology of labeled GCPs in *ex vivo* slices of *Cdh2*^*fl/fl*^*;NeuroD1-Cre* cerebellum was similar to control (Fig. 4 A, B), the median distance of migration, calculated by measuring the distance of the cell soma to the parallel fiber axons, was reduced compared with controls (Control: 120 μm, cKO: 75 μm; *P* < 0.001), indicating a slowed migration rate.

**Fig. 4.**
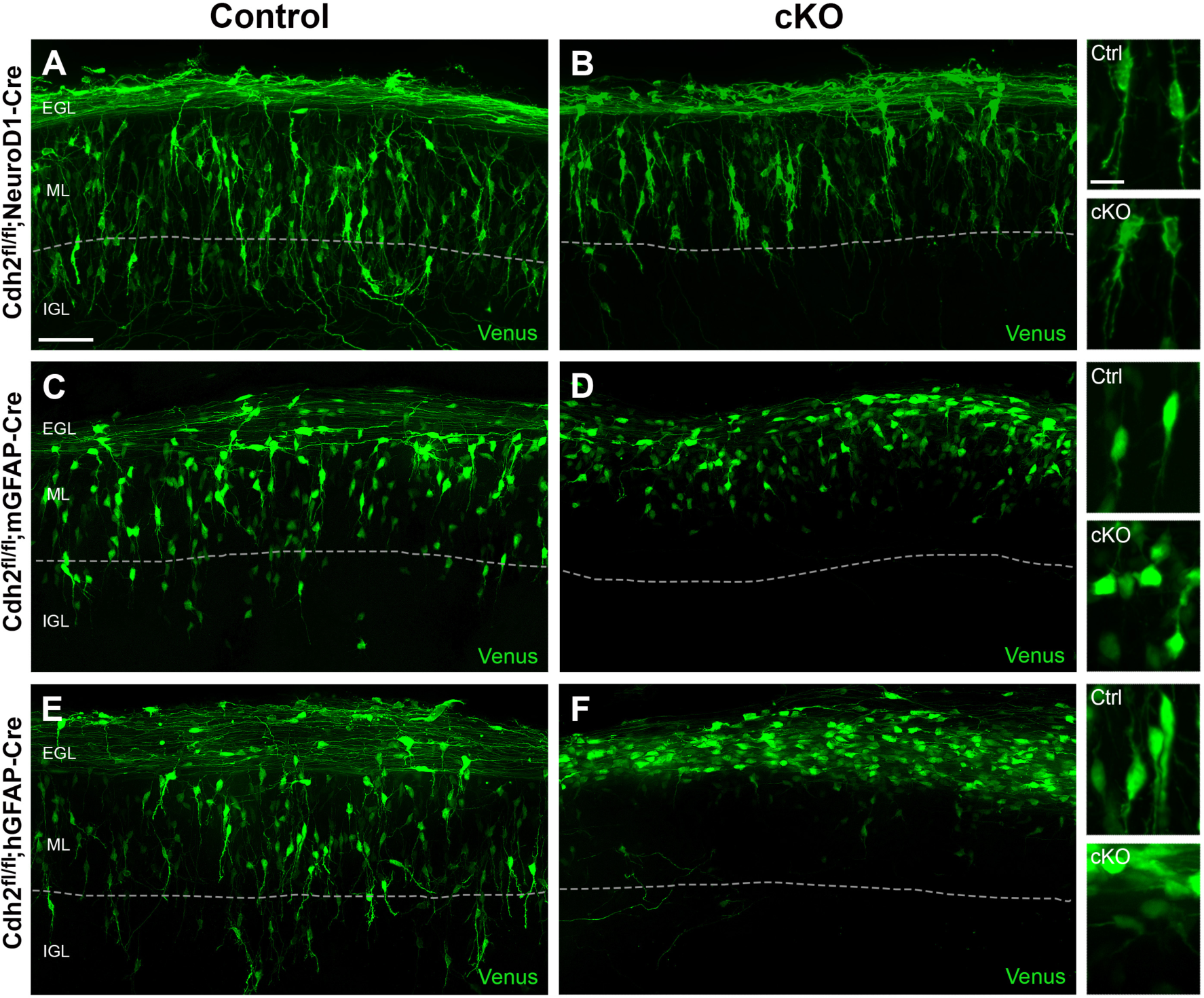
Glial CDH2 is essential for glial-guided neuronal migration. Organotypic *ex vivo* slice cultures prepared from the cerebellum of P8 *Cdh2*^*fl/fl*^ control and *Cdh2* cKO mice, electroporated with *Venus*. In slice cultures from control mice **(A, C, E)**, Venus-expressing GCPs migrated radially across the ML and extended a leading process in the direction of migration. Although most Venus-positive GCPs lacking *Cdh2* **(B)** extended a leading process, their overall migration distance was reduced. In *ex vivo* slices where BG lacked *Cdh2* **(D)**, or where both GCPs and BG lacked *Cdh2* **(F)**, Venus-positive GCPs had a rounded or multipolar morphology, failed to extend a leading process and migrated a shorter distance away from the field of labeled parallel fibers into the ML, indicating a stalled migration. Representative cell morphologies are shown in the insets. Dotted line: ML/IGL boundary. Scale bar represents 50 μm in (A - F) and 10 μm in insets.

While Venus-positive GCPs in *ex vivo* organotypic cultures of both *Cdh2*^*fl/fl*^*;mGFAP-Cre* and *Cdh2*^*fl/fl*^*;hGFAP-Cre* mice appeared to extend parallel fiber axons normally, the GCPs had dramatic morphological defects, as nearly all of the cells were rounded or irregularly shaped, rather than elongated, and failed to extend a leading process in the direction of migration (Fig. 4 C - F). Importantly, virtually all of the Venus-positive cells were stalled in the EGL or the upper portion of the ML, indicating a failure of glial-guided GCP migration.

### Functional Interaction of CDH2 and ASTN1 During Granule Cell Migration

Since glial, but not neuronal, loss of *Cdh2* stalled GCP migration, we hypothesized that ASTN1 may promote glial-guided migration in the absence of neuronal CDH2. To examine the function of ASTN1 in GCPs positive or negative for CDH2, we electroporated the *Venus, Astn1-FL-Venus* or *Astn1-ΔCTD-Venus* plasmids into cerebella of P8 control and *Cdh2*^*fl/fl*^*;NeuroD1-Cre* mice prior to generating organotypic cultures. No significant differences in migration distance or proportion of migrating cells were observed between slice cultures with GCPs expressing *Venus* and *Astn1-FL-Venus* (Fig. 5 A, B, D, E), indicating that ASTN1-FL overexpression did not alter glial-guided migration. However, expression of the *Astn1-ΔCTD-Venus* plasmid in GCPs in control slices reduced the median migration distance by 35 % compared to control slices expressing *Venus* (78 μm and 120 μm, respectively; *P* < 0.001; Fig. 5 C, G), which is a similar reduction to that of the *Cdh2*^*fl/fl*^*;NeuroD1-Cre* slices expressing *Venus* (37 %; Fig. 5 G). This indicates that the ASTN1-ΔCTD protein acted as a dominant negative variant of ASTN1-FL. Expression of *Astn1-ΔCTD-Venus* in GCPs lacking *Cdh2 (Cdh2*^*fl/fl*^*;NeuroD1-Cre)* further reduced migration distance by 39 % compared to the control with *Astn1-ΔCTD-Venus* (48 μm and 78 μm, respectively; *P* < 0.001), and by 60 % compared to the control with *Venus* (48 μm and 120 μm, respectively; *P* < 0.001; Fig. 5 F, G). Importantly, combined disruption of CDH2 and ASTN1 in GCPs resulted in a significant failure to migrate out of the EGL. In addition, expression of *Astn1-ΔCTD-Venus* in GCPs generated a lower proportion of migrating cells, as characterized by their morphology (rounded/multipolar vs. bipolar; Fig. 5 H). These experiments show that GCPs that expressed ASTN1 lacking the domains that bind CDH2 failed to extend a leading process and to migrate. Thus, although homophilic CDH2:CDH2 interactions may contribute to neuron-glial binding, these data demonstrate that heterophilic ASTN1:CDH2 binding is required for glial-guided neuronal migration.

**Fig. 5.**
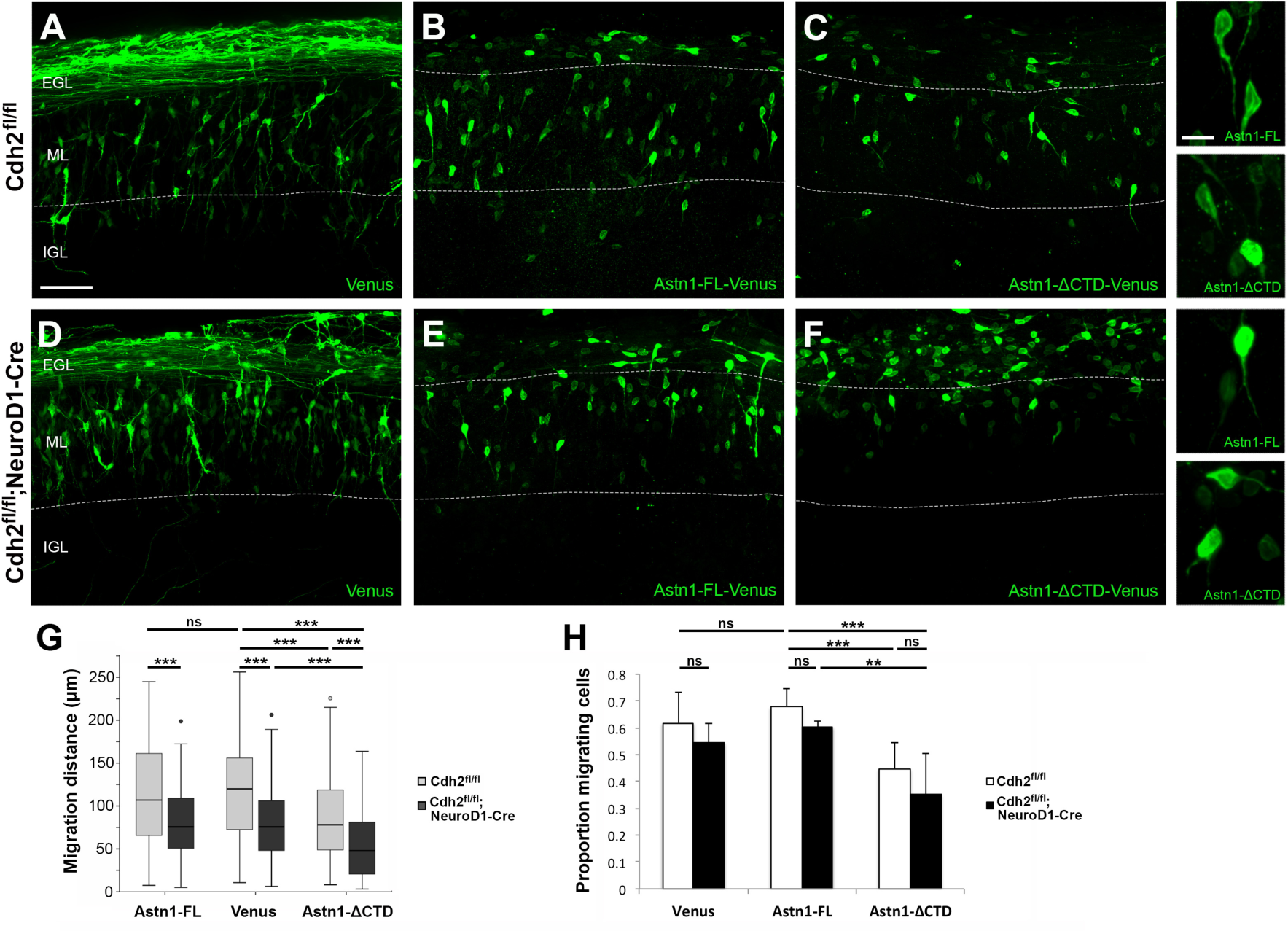
ASTN1 and CDH2 functionally interact to regulate migration. Organotypic slice cultures from the cerebellum of P8 *Cdh2*^*fl/fl*^ and *Cdh2*^*fl/fl*^ *;NeuroD1-Cre* mice, electroporated with *Venus* **(A, D)**, *Astnl-FL-Venus* **(B, E)** or *Astn1-ΔCTD-Venus* **(C, F)**. ASTN1-Venus fluorescence labeled the cell soma and processes but not the parallel fibers. After 60 h, GCPs expressing *Astn1-FL-Venus* migrated a similar distance to that of GCPs expressing *Venus* **(A, B, G)**, also in the absence of neuronal *Cdh2* **(D, E,G)**. However, in slice cultures of control mice where GCPs expressed *Astn1-ΔCTD-Venus*, GCPs migrated a 35 % shorter distance (78 μm) compared to GCPs expressing *Venus* (120 μm) **(C, G)**. Loss of *Cdh2* combined with overexpression of *Astn1-ΔCTD-Venus* in GCPs resulted in more severe migration defects, indicated by a 60 % reduction in migration distance (48 μm) and a higher number of cells stalled in the EGL **(F, G)**. A significantly higher proportion of cells expressing *Astn1-ΔCTD-Venus* were rounded or multipolar **(H)**. Dotted lines: EGL/ML and ML/IGL boundaries. ** *P* < 0.01; *** *P* < 0.001; ns: not significant. Scale bar represents 50 μm in (A - F) and 10 μm in insets.

### Neuron-Glia Attachment is Dependent on Glial Expression of CDH2

To directly analyze the formation of the migration junction in GCPs and BG lacking *Cdh2*, we generated *in vitro* chimeras (23, 31). For these experiments, we purified GCPs or BG using a step gradient of Percoll (30) and mixed and matched GCPs and BG from *Cdh2*^*fl/fl*^ control and *Cdh2*^*fl/fl*^*;hGFAP-Cre* (GCP + BG cKO) cerebella. Purified GCPs from each genotype were electroporated with either a *Venus* plasmid, to visualize the GCP soma and processes, or an *Astn1-FL-Venus* plasmid, to examine the ASTN1 protein localization, and co-cultured with purified glia from each genotype. In control cultures of wild type GCPs and BG, Venus or ASTN1-FL-Venus expressing GCPs adhered to GFAP-labeled BG fibers, formed an elongated profile along the fiber and extended a leading process in the direction of migration (Fig. 6 A, B). ASTN1-FL-Venus localized to the migration junction and the overexpression did not disrupt neuron-glia attachment or migration. Similar results were observed in co-cultures of GCPs lacking CDH2 with control BG (Fig. 6 C, D). A strikingly different result was seen when either control or CDH2-deficient GCPs were cultured with BG lacking CDH2. In both cases, although the ASTN1 protein localized to the basal portion of the soma, the neurons failed to form a migration junction (Fig. 6 E - H). Measurement of the distance between the GCP soma and BG fiber confirmed this observation (Fig. 6 I). In addition, GCPs were rounded or multipolar, rather than elongated as seen on control BG fibers, and failed to extend a leading process. Thus, formation of the migration junction between GCPs and BG fibers required glial CDH2.

**Fig. 6.**
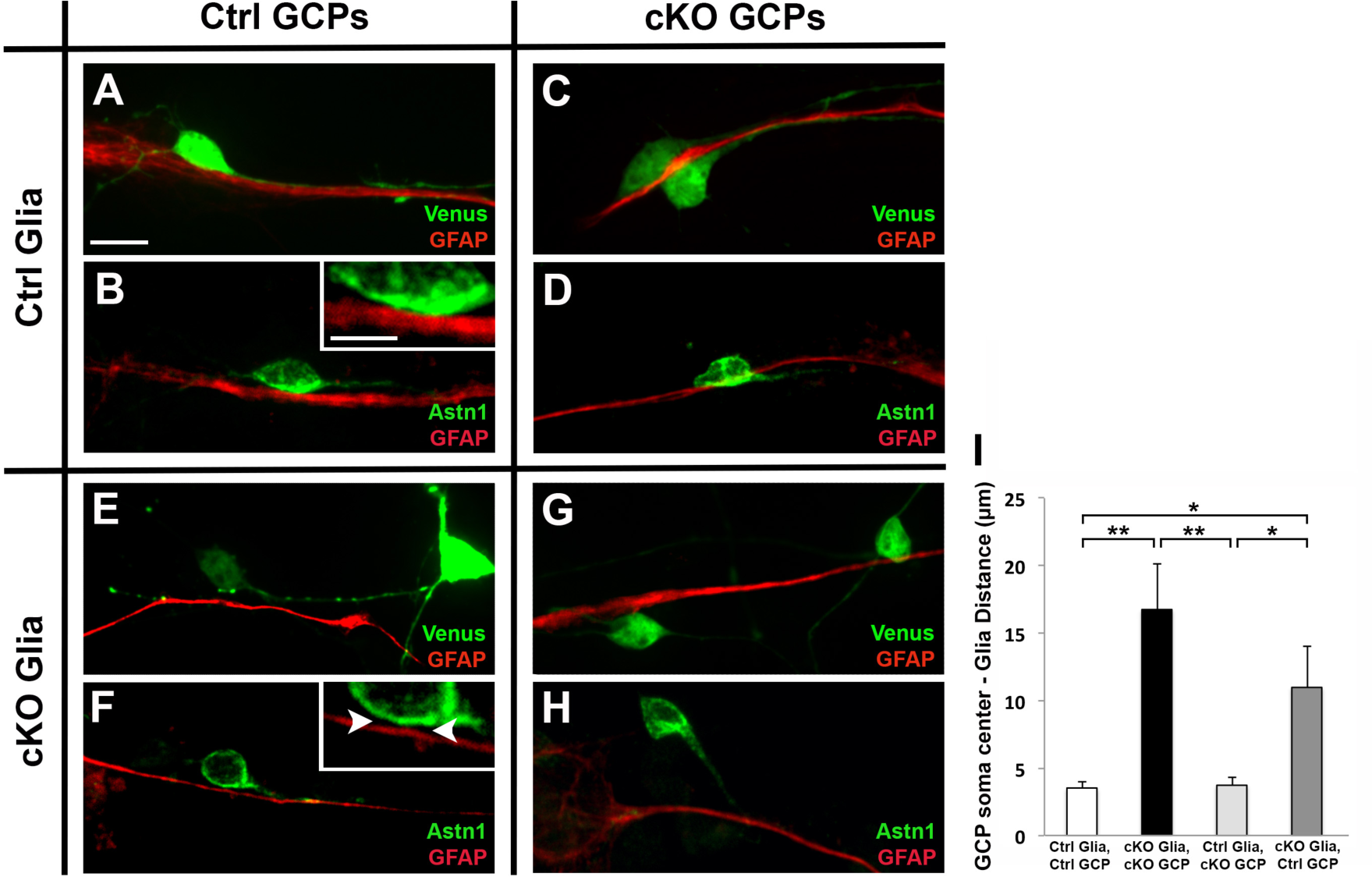
Neuron-glia attachment is dependent on glial CDH2. *In vitro* chimera co-cultures of GCPs and BG purified from *Cdh2*^*fl/fl*^ (control) or *Cdh2*^*fl/fl*^*;hGFAP-Cre* (cKO) mice at P7. GCPs were electroporated with *Venus* **(A, C, E,G)** or *Astn1-Venus* **(B, D, F, H)**. GCPs attached and migrated along BG fibers expressing CDH2 **(A - D)**, irrespective of the GCP genotype. In contrast, GCPs co-cultured with BG lacking *Cdh2* did not form a migration junction with the glial fibers **(E - H)** and had an increased separation between the GCP somas and the glial fibers **(I)**. Note the gap between the neuronal soma and BG fiber even as the GCP process contacts the fiber (inset in F, arrowheads). * *P* < 0.05; ** *P* < 0.01. Scale bar represents 10 μm in (A - H), and 5 μm in insets in (B) and (F).

Taken together, our results propose a bridge model in which a *cis* complex of ASTN1 and CDH2 in the neuronal membrane interacts in *trans* with CDH2 on the glial fiber (Fig. 7) to promote glial-guided migration.

**Fig. 7.**
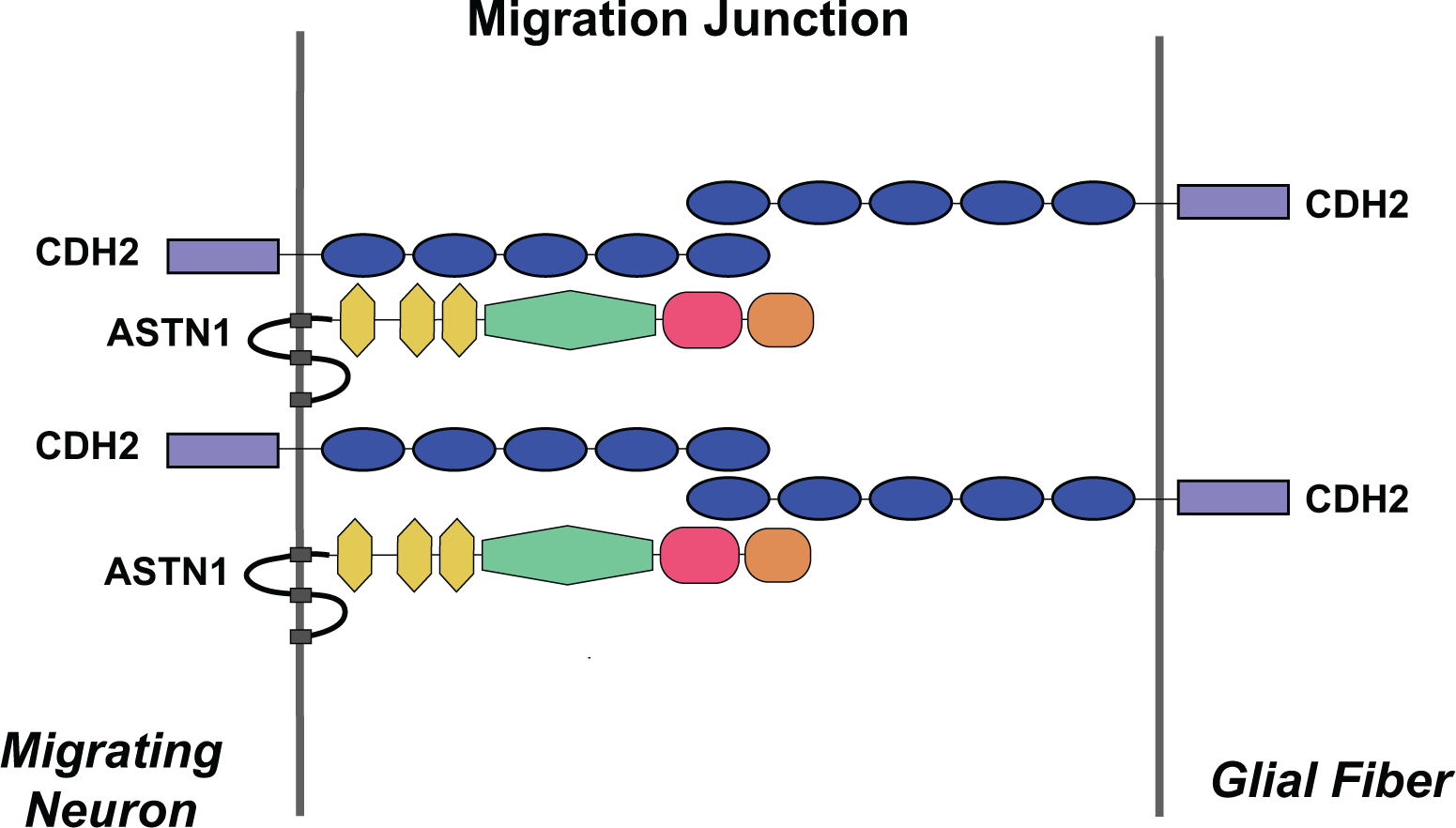
Bridge model of ASTN1 and CDH2 in the migration junction. Proposed model of the ASTN1:CDH2 *cis* and *trans* asymmetric bridge complex in the neuron-glial migration junction. Blue oval: cadherin domain; yellow hexagon: EGF-like domain; green hexagon: MACPF domain; magenta oval: FNIII domain; brown oval: ANX-like domain.

## DISCUSSION

This study reports evidence for the formation of a *cis* and *trans* asymmetric bridge complex between two families of CNS guidance receptors, CDH2 and ASTN1, as well as the discovery that CDH2 is the glial ligand for ASTN1, the neuronal receptor for glial-guided migration. These findings are supported by biochemical evidence that ASTN1 binds CDH2 in *cis*, by S2 adhesion assays showing that ASTN1 binds CDH2 in *trans*, by genetic conditional loss of function studies showing defects in glial-guided migration in mice lacking *Cdh2* in glia, and by *in vitro* chimeras showing a failure of GCPs to form a migration junction with glia lacking CDH2. Overexpression of an ASTN1 variant lacking the binding domain for CDH2 confirmed that homophilic CDH2 binding between GCPs and glia is not sufficient to support neuronal migration. Thus, heterophilic bridge complexes of ASTN1 and CDH2 are required for glial-guided neuronal migration in the developing cerebellum.

Support for *cis* interactions of ASTN1 and CDH2 was provided by immunoprecipitation assays, which further confirmed that three specific extracellular domains of ASTN1 - MACPF, FNIII and ANX-like - were required for ASTN1 binding to CDH2. This finding provided the molecular basis for studies on the overexpression of ASTN1 lacking these domains, which appeared to act as a dominant negative. The biochemical findings were also supported by S2 cell adhesion assays, showing that heterophilic ASTN1:CDH2 binding supports cell-cell adhesion, in addition to homophilic CDH2-induced cell adhesion. Thus, CDH2 binds ASTN1 in both *cis* and *trans* and in intercellular adhesion. These findings are reminiscent of *cis* and *trans* binding between CDH2 and the AMPA receptor subunit GluR2 in dendritic spines (32).

Conditional loss of function experiments have been instrumental in defining the site of action of specific neuronal or glial receptors. The significance of glial CDH2, relative to neuronal CDH2, in glial-guided migration was evident from our finding that a glial loss of *Cdh2* resulted in dramatic effects on GCP morphology and a failure to adhere to glial fibers, whereas a neuronal loss of *Cdh2* did not affect neuron-glia binding or stall migration. We therefore propose that the binding of ASTN1 to glial CDH2 is sufficient to promote neuron-glia attachment and glial-guided migration. The finding that a loss of *Cdh2* in GCPs slowed glial-guided migration suggests that the *cis* interaction of ASTN1 and CDH2 is also important for migration, likely by stabilizing the migration junction between the neuron and the glial fiber. One mechanism for stabilizing the migration junction would be through endosomal recycling of ASTN1 back to the plasma membrane - a role that has previously been ascribed for ASTN2, the second member of the Astrotactin family (7). Indeed, CDH2 has been reported to function in AMPA receptor trafficking by increasing surface expression of AMPA receptors in neurons (33). Interestingly, we detected an interaction of CDH2 with ASTN2 in cerebellar GCPs by co-IP (data not shown), suggesting that CDH2 is involved in the endocytic Astrotactin protein complex.

Other studies have proposed that homophilic CDH2:CDH2 binding regulates neuron-glia attachment and migration (19, 20). Our bridge model does not exclude this scenario; however, the discrepancy in migration defects caused by the cell-specific loss of *Cdh2* in neurons or glia indicates that additional proteins to CDH2 are involved in the complex. Thus, we suggest that homophilic *trans* CDH2 interactions contribute to the formation of a migration junction, but that the CDH2:CDH2 complex is not sufficient for migration. The present study extends the general role for cadherins in homophilic cell-cell interactions by directly demonstrating that both *cis* and *trans* ASTN1:CDH2 interactions function in glial-guided migration.

Further support for the asymmetric bridge complex comes from the finding that a combined deletion of *Cdh2* in both GCPs and BG produced migration defects similar to the glia-specific deletion. While the combined deletion also resulted in developmental defects in the cerebral cortex (Horn & Hatten, unpublished), this was likely due to earlier expression of the hGFAP promoter (E13.5) (29), compared to the mGFAP promoter (postnatal) (28). Moreover, the slightly more disorganized ML/IGL boundary in the cerebellum of the *Cdh2*^*fl/fl*^*;hGFAP-Cre* mice was also likely attributable to the difference in promoter expression. Importantly, all migration assays in this study were comparable as they assessed GCP migration at postnatal stages (P5 - P8) where *Cdh2* was deleted in all three lines (Fig. S2).

Heterophilic interactions between different cadherins have been reported (34, 35). However, no cerebellar defects have been described in mice with a targeted deletion of R-cadherin (36), M-cadherin (37) or cadherin-11 (38), which are expressed in the postnatal mouse cerebellum (39), suggesting that these cadherins do not function in GCP migration. Still, we cannot exclude that a partial compensation of other cadherins occurred in the neuron-specific *Cdh2* cKO. Interestingly, cadherin-11 was recently shown to regulate neural crest migration via binding of its cleaved EC1-3 domains to ErbB2 (40). Moreover, CDH2 has been demonstrated to interact in *trans* with the AMPA receptor subunit GluR2 to regulate spine formation (32). This corroborates our findings that cell adhesion and migration can be regulated by cadherins independently of homophilic or compensatory cadherin bindings.

The heterophilic ASTN1:CDH2 interaction was shown to occur via the C-terminal ectodomain of ASTN1, which included the MACPF, FNIII and ANX-like domains (Fig. 1 B). We were unable to pinpoint whether a single domain binds CDH2, since ASTN1 constructs lacking only the MACPF or FNIII domains failed to localize to the cell surface. Expression of ASTN1-ΔCTD in control organotypic slices did not fully stall GCP migration, but significantly slowed migration similar to the neuronal loss of *Cdh2*. Slowed GCP migration is consistent with previous migration studies on *Astn1* mutant mice (13). However, it is possible that endogenous ASTN1 may not have been fully competed out by ASTN1-ΔCTD and still contributed to GCP migration. In addition, we cannot exclude that other adhesion proteins are also involved in the ASTN1:CDH2 bridge complex. Nevertheless, the combined deletion of *Cdh2* with ASTN1-ΔCTD expression in GCPs resulted in a migration failure similar to the glial *Cdh2* cKO, demonstrating that a *cis* interaction of ASTN1 and CDH2 in GCPs promotes migration. This is consistent with the observation that CDH2 acts in combination with nectin-based adhesion to regulate radial glia-independent somal translocation (41).

Although the intracellular aspect of ASTN1 does not contain any domains known to be involved in intracellular signaling pathways, it is possible that the complex of ASTN1:CDH2 functions in intracellular signaling during migration. The best characterized signaling cascades involving CDH2 are beta-catenin (42) and GTPases (22, 43). Efforts to examine a role for active beta-catenin in GCP migration were complicated by the fact that beta-catenin appears to function in both parallel fiber extension and migration, and the fact that we did not detect changes in active beta-catenin signaling in migrating GCPs of the *Cdh2* mutants (Horn & Hatten, unpublished). While recent studies support a key role for Rho, Rab and Rap GTPases in the control of CDH2 function (18, 22, 43) and in GCP migration via actin-regulatory pathways (10), their signaling role in CDH2:ASTN1 complexes remains to be determined. In addition, since prior EM studies revealed the presence of microfilaments in the migration junction (6) where ASTN1 localizes (11), it will be important to assay whether cytoskeletal elements, including actin-binding proteins, localize to the migration junction.

Our study provides the first direct demonstration of *cis* and *trans* interactions of CDH2 with a CNS migration receptor, and raises the possibility for a general function for heterophilic cadherin-receptor complexes in the formation of a wide range of cell-cell junctions.

## MATERIALS AND METHODS

### Animals

B6.129S6(SJL) *Cdh2*^*fl/fl*^ mice (backcrossed to C57Bl/6) carrying loxP sites flanking exon 1 of the *Cdh2* gene (Jackson Laboratory, Stock # 007611) were crossed with *Tg(NeuroD1-Cre) RZ24, Tg(hGFAP-Cre) PK90* (both provided by Dr. Nathaniel Heintz/Gensat) or *Tg(mGFAP-Cre)* (provided by Dr. Alexandra Joyner) lines. *Cdh2p^fl/+^;NeuroD1-Cre, Cdh2p^fl/+^;mGFAP-Cre* and *Cdh2*^*fl/+*^*;hGFAP-Cre* progeny were crossed with *Cdh2*^*fl/fl*^ mice to generate *Cdh2^fl/fl^;Cre^+/-^* experimental mice and *Cre-* negative *Cdh2*^*fl/fl*^ control littermates. Genotyping details are described in SI Methods. All procedures were performed according to the guidelines approved by the Rockefeller University Institutional Animal Care and Use Committee.

### DNA constructs

See SI Methods for details.

### Granule cell/Bergmann glia co-cultures

Co-cultures of granule cells and Bergmann glia from P7 cerebella were prepared as described previously (30). Briefly, dissociated cerebellar cell suspension was applied to a two-step gradient of 35 %/60 % Percoll (Sigma-Aldrich) in Tyrode’s solution containing 2 mM EDTA. The Percoll gradient was centrifuged at 3000 rpm for 10 minutes at 4 ^o^C. The large cell fraction at the interface of the Tyrode’s solution and 35 % Percoll (Bergmann glia) and the small cell fraction at the interface between the layer of 35 % Percoll and 60 % Percoll (granule cells) were subsequently washed and preplated in granule cell medium (see SI Methods) on untreated petri dishes at 35 ^o^C/5 % CO_2_ for 20 minutes to remove fibroblasts. For the glial cell fraction, the unbound cell suspension was thereafter cultured on 0.1 mg/ml poly D-lysine (Millipore) pre-coated 12 mm coverslips for 1 h at 35 ^o^C/5 % CO_2_. The unbound cell suspension was then removed and the remaining glial cells, which attached well to the coated coverslips, were cultured in granule cell medium at 35 ^o^C/5 % CO_2_. For the granule cell fraction, the cell suspension was transferred from the petri dish to a 60 mm tissue culture dish and incubated for 30 minutes at 35 ^o^C/5 % CO_2_. The dish was then tapped to dislodge the granule cells and the cell suspension was transferred to a new tissue culture dish and the process repeated for maximal purification of granule cells. Purified granule cells were then electroporated with an Amaxa Mouse Neuron Nucleofection kit (Lonza, Cologne, Germany) as described in SI Methods. Granule cells were added to the Bergmann glia cell cultures at a ratio of 5:1 to the number of glial cells. The total numbers were ~750,000 granule cells and ~150,000 glial cells per 12 mm coverslip. The co-cultures were incubated at 35 ^o^C/5 % CO_2_ for 48 - 72 h before analyzing neuron-glial attachment and granule cell migration.

### Organotypic slice cultures

The method is described in previous work (10) and detailed in SI Methods. Briefly, P8 cerebella from *Cdh2*^*fl/fl*^ and *Cdh2* cKO littermates were dissected out and electroporated with *pCIG2-Venus* (0.5 μg/μl), *pCIG2-Astn1-FL-Venus* (1 μg/μl) or *pCIG2-Astn1-ΔCTD-Venus* (1 μg/μl) plasmids dorsal to ventral for 50 ms at 80 V, for a total of 5 pulses with an interval of 500 ms between pulses, using an *electro-square-porator, ECM* 830 (BTX Genetronics). The cerebella were then embedded in 3 % agarose in HBSS, and 250 μm horizontal slices were made using a Leica VT1000S vibratome. Slices were placed on MilliCell CM 0.4 μm culture plate inserts (Millipore) and cultured for 60 hours at 35 ^o^C/5% CO_2_.

### Immunohisto/cytochemistry

Brains were dissected out and fixed in 4 % paraformaldehyde in PBS at 4 ^o^C overnight and thereafter cryoprotected in 20 % sucrose in PBS at 4 ^o^C overnight. Sagittal cryosections (25 μm), organotypic slice cultures or granule cell/Bergmann glia co-cultures were processed for immunohistochemistry as described in SI Methods.

### BrdU labeling

50 μg of BrdU in PBS (BD Biosciences) per gram of body weight was injected subcutaneously in the neck of P5 *Cdh2*^*fl/fl*^ and *Cdh2* cKO littermates. The brains were dissected out 48 hours later and processed for immunohistochemistry as described in SI Methods.

### Immunoprecipitation and Western blotting

Briefly, transfected HEK 293T cells (clone 17; ATCC CRL-11268) or whole cerebella were extracted in ice-cold lysis buffer (see SI Methods) and precleared with 25 μl Protein G/A Agarose beads (Calbiochem). The lysates were incubated with 3 μg of a rabbit GFP antibody (Invitrogen), rabbit Astn1 antibody (12), or normal rabbit IgG (Santa Cruz Biotechnology) for 2 hours at 4 ^o^C. Immunoprecipitates were collected on 50 μl Protein G/A Agarose beads by overnight rotation at 4 ^o^C, washed with lysis buffer and resuspended in 50 μl 2X Laemmli buffer. Western blotting was performed as described in SI Methods.

### S2 cell adhesion assay

*Drosophila* Schneider 2 (S2) cells (Life Technologies) were transfected for 24 hours as described in SI Methods, and 1.5×10^6^ cells from each condition were mixed together at a density of 3×10^6^ cells/well (1×10^6^ cells/ml) and shaken gently at 28 ^o^C for up to 2 hours to allow aggregation. Cells were imaged on a Carl Zeiss Axiovert 135 fluorescent microscope with a 20X objective immediately after the conditions were set up (T = 0) and after 30 minutes, 1 h and 2 h. Cells expressing GFP (CDH2) or mCherry (control or ASTN1) were quantified in each aggregate and the proportion of mCherry-positive cells per aggregate was calculated for each condition and time point. For full details, see SI Methods.

### Flow cytometry

Transfected HEK 293T cells were harvested in 1 mM EDTA in PBS. The surface fraction of Venus-linked ASTN1 variants was labeled with rabbit anti-GFP (1:5,000; Invitrogen) for 20 minutes at 4 ^o^C followed by Alexa-647 donkey anti-rabbit (1:5,000; Life Technologies) for 25 minutes at 4 ^o^C. Flow cytometry analysis on BD Accuri C6 (BD Biosciences) was carried out using 488 nm and 640 nm lasers as described in SI Methods.

### Statistical analyses

See SI Methods for details. Differences between conditions were determined using unpaired *t*-tests for equal or unequal variances, except for migration distance in the slice cultures where Kruskal-Wallis and Mann-Whitney U non-parametric tests were used. Significance was set at *P* < 0.05 (two-sided). In the bar diagrams, data are presented as means with error bars representing the standard deviations. The migration distance data from the slice cultures are presented in box plots.

## ACKNOWLEDGMENTS

We are grateful to Dr. Eve-Ellen Govek for advice on carrying out organotypic slice culture assays and for comments on the manuscript, to Dr. Nathaniel Heintz and GENSAT for providing the *NeuroD1-Cre* and *hGFAP-Cre* lines, to Dr. Alexandra Joyner for providing the *mGFAP-Cre* line as well as for helpful discussions, to Dr. Richard Huganir for providing the *pRK5-Cdh2* and *pRK5-Cdh2-A390* constructs, to Dr. Leslie Vosshall for providing the *pAc5-STABLE2* construct, to Dr. Franck Polleux for the *pCIG2* construct, and to Drs. David Solecki and Robert Gilbert for valuable comments on the manuscript. Cell surface protein measurements were carried out in the Rockefeller University Flow Cytometry Resource Center, for which we thank Drs. Svetlana Mazel and Stanka Semova for assistance. Bright-field and confocal microscopy was partly carried out in the Rockefeller University Bio-Imaging Resource Center, for which we thank Drs. Alison North, Christina Pyrgaki and Tao Tong for assistance. This study was supported by funding from the Swedish Research Council (Z.H.), the Nicholson Postdoctoral Exchange Program between the Rockefeller University and Karolinska Institutet (Z.H.), and the Hoffmann Trust (M.E.H., H.B.).

## SUPPLEMENTARY INFORMATION

### SI Methods

#### Animals

B6.129S6(SJL) *Cdh2*^*fl/fl*^ mice (backcrossed to C57Bl/6) carrying loxP sites flanking exon 1 of the *Cdh2* gene (Jackson Laboratory, Stock # 007611) were crossed with *Tg(NeuroD1-Cre) RZ24, Tg(hGFAP-Cre) PK90* (both provided by Dr. Nathaniel Heintz/Gensat) or *Tg(mGFAP-Cre)* (provided by Dr. Alexandra Joyner) lines. The NeuroD1 promoter commences at E16 in postmitotic granule cells (27), the mGFAP promoter after P0 in Bergmann glial cells (28), and the hGFAP promoter at E13.5 in neural progenitors (29). *Cdh2^fl/+^;NeuroD1-Cre, Cdh2^fl/+^;mGFAP-Cre* and *Cdh2*^*fl/+*^*;hGFAP-Cre* progeny were crossed back to *Cdh2*^*fl/fl*^ mice. The *Cdh2^fl/fl^;Cre^+/-^* experimental mice were only compared with Cre-negative *Cdh2*^*fl/fl*^ control littermates. Genotyping of the *Cdh2*^*fl/fl*^ allele was carried out by PCR using 5’-CCA AAG CTG AGT GTG ACT TG and 5’-TAC AAG TTT GGG TGA CAA GC primers, and of the *Cre* allele using 5’-GGA CAT GTT CAG GGA TCG CCA GGC G and 5’-GCC AGA TTA CGT ATA TCC TGG CAG CG primers. The number of animals for each phenotypic analysis is stated in the Results.

All animal work was performed as required by the United States Animal Welfare Act and the National Institutes of Health’s policy to ensure proper care and use of laboratory animals for research, and under established guidelines and supervision by the Institutional Animal Care and Use Committee (IACUC) of The Rockefeller University. Mice were housed in accredited facilities of the Association for Assessment of Laboratory Animal Care (AALAC) in accordance with the National Institutes of Health guidelines.

#### DNA constructs

*Venus* cDNA was PCR generated from *pMSCXβ-Venus* (8) template with the following primers: Venus forward (EcoRI) primer 5’-GAG AAG <u>GAA TTC</u> ACC *ATG* GTG AGC AAG GGC GAG GAG and Venus reverse (NotI) primer 5’-GAG AAG <u>GCG GCC GC</u>T *TAC* TTG TAC AGC TCG TCC ATG CCG (restriction sites underlined and start/stop codons in italics). The PCR product was inserted into*pCIG2* plasmid (kindly provided by Dr. Franck Polleux) digested with EcoRI and NotI. The *pCIG2-Astn1-Venus* plasmid was generated by fusing the *Venus* sequence in frame with the 3’ end of the *Astn1* coding sequence by joining PCR. The resulting *Astn1-Venus* fusion insert (EcoRI/NotI) was subcloned, along with the *Astn1* cDNA fragment (XmaI/EcoRI), into the *pCIG2* vector. Finally, the 5’ region (231 bp) of the *Astn1* cDNA was inserted between the XhoI and XmaI sites. For *pCIG2-Astn1-ΔCTD-Venus*, the *pCIG2-Astn1-Venus* plasmid was digested with PstI and the larger fragment was purified. Thereafter, a gBlock DNA fragment (Integrated DNA Technologies, Coralville, IA), flanked by PstI sites and containing a fusion between *Astn1* coding sequence base pair number 2160 and the *Venus* start site, was inserted, which generated a 1749 bp deletion of the *Astn1* 3’ region resulting in a 583 amino acid deletion in the ASTN1 C-terminus. The *pRK5-Cdh2-Myc* and *pRK5-Cdh2-Δ390-Myc* plasmids were kindly provided by Dr. Richard Huganir. The *pAc5-STABLE2-Neo* plasmid containing the *Drosophila* Actin5C promoter (25) was kindly provided by Dr. Leslie Vosshall. To generate the *pAc5-Astn1-mCherry* plasmid, *Astn1* cDNA was amplified by PCR with the following primers: Astn1 forward (XbaI) primer 5’-CTT <u>TCT AGA</u> *ATG* GCT TTA GCC GGG CTC TG and Astn1 reverse (XhoI) primer 5’-CTT <u>CTC GAG</u> *CTA* GAT GTC TTT GCT GTC CC (restriction sites underlined and start/stop codon in italics). The PCR product was inserted between the XbaI and XhoI sites in the *pAc5-STABLE2-Neo* plasmid. The *pAc5-Cdh2-GFP* and *pAc5-Cdh2-A390-GFP* plasmids were generated by amplifying the *Cdh2* and *Cdh2-Δ390* cDNA with the primers: Cdh2 forward (EcoRI) 5’-GC<u>G AAT TC</u>*A TGT* GCC GGA TAG CGG GA and Cdh2 reverse (NotI) 5’-TT<u>G CGG CCG C</u>TG TCG TCA CCA CCG CC (restriction sites underlined and start codon in italics). The *Cdh2* and *Cdh2-Δ390* PCR products were inserted into the *pAc5-STABLE2-Neo* plasmid digested with EcoRI and NotI. The *pAc5-STABLE2-Neo* plasmid contains T2A sequences to allow bicistronic expression of GFP and mCherry instead of protein fusions.

#### Granule cell/Bergmann glia co-cultures

Cerebella were dissected out from P7 mice and the meninges were carefully removed. The cerebella were then incubated in Trypsin-DNase for 5 minutes at 37 ^o^C followed by gentle trituration in DNase to dissociate the tissue into single cells. The cell suspension was filtered through a 40 μm nylon mesh filter (BD Biosciences) and then applied to a two-step gradient of 35 %/60 % Percoll (Sigma-Aldrich) in Tyrode’s solution containing 2 mM EDTA. The Percoll gradient was centrifuged at 3000 rpm for 10 minutes at 4 ^o^C. The large cell fraction at the interface of the Tyrode’s solution and 35 % Percoll (Bergmann glia) and the small cell fraction at the interface between the layer of 35 % Percoll and 60 % Percoll (granule cells) were transferred to Tyrode’s solution and centrifuged at 2000 rpm for 5 minutes at 4 ^o^C. The supernatant was removed and the cell pellets were resuspended in granule cell medium [Basal Medium Eagle (BME; Gibco, cat # 21010) supplemented with 30 % D-glucose (Sigma-Aldrich), 10 % horse serum (Gibco), 2 mM L-glutamine (Gibco) and 100 U/ml penicillin/streptomycin (Gibco)]. The cell suspensions were transferred to untreated petri dishes and incubated at 35 ^o^C/5 % CO2 for 20 minutes to remove fibroblasts.

For the glial cell fraction, the unbound cell suspension was removed and cultured on 0.1 mg/ml poly D-lysine (Millipore) pre-coated 12 mm coverslips for 1 h at 35 ^o^C/5 % CO_2_. The plate was then tapped to dislodge granule cells that had accompanied the glial cell fraction, and the unbound cell suspension was removed. The remaining glial cells, which attached well to the coated coverslips, were cultured in granule cell medium at 35 ^o^C/5 % CO_2_. For the granule cell fraction, the cell suspension was transferred from the petri dish to a 60 mm tissue culture dish and incubated for 30 minutes at 35 ^o^C/5 % CO_2_. The dish was then tapped to dislodge the granule cells and the cell suspension was transferred to a new tissue culture dish and the process repeated for maximal purification of granule cells. The purified granule cells were then electroporated with an Amaxa Mouse Neuron Nucleofection kit (Lonza, Cologne, Germany), using 3×10^6^ cells, 6 μg *pCIG2-Venus* DNA or 15 μg *pCIG2-Astn1-Venus* DNA, and the O-005 setting in the Amaxa Nucleofector II. Immediately after electroporation, granule cells were recovered in granule cell medium at 35 ^o^C/5 % CO_2_ for 15 minutes, centrifuged at 2000 rpm for 5 minutes at 4 ^o^C, resuspended in granule cell medium and subsequently added to the Bergmann glia cell cultures at a ratio of 5:1 to the number of glial cells. The total numbers were ~750,000 granule cells and ~150,000 glial cells per 12 mm coverslip. The co-cultures were incubated at 35 ^o^C/5 % CO_2_ for 48 - 72 h before analyzing neuron-glial attachment and granule cell migration along glial fibers.

#### Organotypic slice cultures

P8 cerebella from *Cdh2*^*fl/fl*^ and *Cdh2* cKO littermates were dissected out in Hank’s Balanced Salt Solution (HBSS) containing 2.5 mM HEPES (pH 7.4), 46 mM D-glucose, 1 mM CaCl_2_, 1 mM MgSO_4_, 4 mM NaHCO_3_, and Phenol Red on ice. The dissection medium was then removed and *pCIG2-Venus* DNA was diluted to 0.5 μg/μl and *pCIG2-Astn1-FL-Venus* or *pCIG2-Astn1-ΔCTD-Venus* DNA diluted to 1 μg/μl in HBSS. The cerebella were soaked in the DNA for 15 minutes on ice, and were then transferred one at a time into the well of an electroporation chamber (Protech International Inc. CUY520P5 platinum electrode L8×W5×H3 mm, 5 mm gap) that was placed on ice. The cerebella were electroporated dorsal to ventral for 50 ms at 80 V, for a total of 5 pulses with an interval of 500 ms between pulses, using an *electro-square-porator, ECM* 830 (BTX Genetronics). The cerebella were then removed from the chamber and placed on ice to recover for 10 minutes.

Subsequent to electroporation, the cerebella were embedded in 3 % agarose in HBSS, and 250 μm horizontal slices were made using a Leica VT1000S vibratome set at a speed of 3 and frequency of 6. Slices were then placed on MilliCell CM 0.4 μm culture plate inserts (Millipore) in a 6 well plate with 1.5 ml of culture medium [BME (Gibco), 25 mM D-glucose (Sigma-Aldrich), 2 mM L-glutamine (Gibco), 1X insulin/transferrin/selenium (Sigma-Aldrich), 100 U/ml penicillin/streptomycin (Gibco)] below the insert. The organotypic slices were incubated at 35 ^o^C/5 % CO2 for 60 hours before fixing with 4 % paraformaldehyde (PFA)/4 % sucrose in phosphate-buffered saline (PBS) for 2 hours at room temperature.

For immunostaining, the agarose was carefully removed from the slices and the slices were permeabilized and blocked overnight at 4 ^o^C in 10 % normal donkey serum (NDS)/0.3 % Triton X-100 in PBS. The slices were then incubated overnight at 4 ^o^C with a rabbit anti-GFP antibody (1:2000; Invitrogen) diluted in 10 % NDS/0.3 % Triton X-100 in PBS, followed by washes in 10 % NDS/0.3 % Triton X-100 in PBS and overnight incubation at 4 ^o^C with an Alexa Fluor 488 conjugated secondary antibody (Life Technologies) diluted in 10 % NDS/0.3 % Triton X-100 in PBS. The following day, the slices were washed in PBS and mounted with ProLong Gold anti-fade reagent (Molecular Probes). The slices were imaged with a Carl Zeiss Axiovert 200M/ Perkin Elmer Ultraview spinning disk confocal microscope equipped with a 25X objective.

#### Nissl staining

Brains were dissected out and fixed in 4 % PFA in PBS at 4 ^o^C overnight and thereafter cryoprotected in 20 % sucrose in PBS at 4 ^o^C overnight. The brains were then embedded in Tissue-Tek O.C.T. compound (Sakura Finetek, Torrance, CA) and rapidly frozen. 25 μm sagittal sections were prepared using a Leica CM 3050S cryostat. Sections were postfixed in 4 % PFA in PBS for 15 minutes, rinsed in PBS and incubated in 0.1 % cresyl violet for 5 minutes. The sections were then rinsed in water and dehydrated in a dilution series of ethanol (50, 70, 95, 100 %) and Xylene before mounting the slides in Eukitt mounting medium (Sigma-Aldrich). The sections were imaged on a Carl Zeiss Axioplan 2 microscope with a 5X objective.

#### Immunohisto/cytochemistry

Brains were prepared as described above. Sagittal cryosections (25 μm) or granule cell/Bergmann glia co-cultures were fixed in 4 % PFA in PBS for 15 min, rinsed in PBS and blocked/permeabilized in 5 % NDS/0.3 % Triton X-100 in PBS for 45 minutes. Primary antibodies were incubated overnight at 4 ^o^C, followed by PBS washes and incubation with Alexa Fluor 488 or 555 conjugated secondary antibodies (1:400; Life Technologies) for 2 h at room temperature. After subsequent PBS washes, sections were mounted in Vectashield mounting medium (Vector Laboratories, Burlingame, CA) and cell cultures were mounted in ProLong Gold anti-fade reagent (Molecular Probes). The primary antibodies used were mouse anti-N-cadherin (1:500; BD Transduction Laboratories), rabbit anti-Astn1 (1:200) (12), rabbit anti-BLBP (1:300; Millipore), mouse anti-GFAP (1:500; Sigma-Aldrich), mouse anti-NeuN (1:200; Chemicon), sheep anti-BrdU (1:100; Abcam), rabbit anti-active Caspase-3 (1:400; Cell Signaling Technology), mouse anti-phospho-histone H3 (1:100; Cell Signaling Technology), mouse anti-Calbindin D28-k (1:500; Swant) and rabbit anti-GFP (1:1000; Invitrogen). Primary antibodies were titrated to determine the optimal dilutions, and negative control labelings were included with the respective primary antibody omitted. The slides were imaged on a Carl Zeiss Axiovert 200M/ Perkin Elmer Ultraview spinning disk confocal microscope or a Leica DMI 6000 (TCS SP8) confocal microscope with 25X and 40X objectives.

#### BrdU labeling

50 μg of BrdU in PBS (BD Biosciences) per gram of body weight was injected subcutaneously in the neck of P5 *Cdh2* ^*fl/fl*^ and *Cdh2* cKO littermates. The brains were dissected out 48 h later and fixed in 4 % PFA in PBS at 4 ^o^C overnight, followed by cryoprotection in 20 % sucrose in PBS at 4 ^o^C overnight. The brains were then embedded in Tissue-Tek O.C.T. compound (Sakura Finetek, Torrance, CA) and rapidly frozen. 25 μm sagittal sections were prepared using a Leica CM 3050S cryostat. The sections were postfixed in 4 % PFA in PBS for 15 min, washed in PBS, and then incubated in 2N HCl for 30 minutes at 37°C followed by 0.1 M Sodium Borate, pH 8.5, for 10 minutes at room temperature. The sections were then processed for immunohistochemistry as described above.

#### Cell culture

Human embryonic kidney (HEK) 293T cells (clone 17; ATCC CRL-11268) were cultured at 37 ^o^C/5 % CO_2_ in Dulbecco’s modified Eagle’s medium (Gibco, cat # 11995) supplemented with 10 % fetal bovine serum (FBS; Gibco), 2 mM L-glutamine (Gibco) and 100 U/ml penicillin/streptomycin (Gibco). Cells at 50–60 % confluency were transfected with 4 μg DNA constructs using Lipofectamine 2000 (Invitrogen) according to the manufacturer’s instructions.

*Drosophila* Schneider 2 (S2) cells (Life Technologies) were cultured at 28 ^o^C in Schneider’s *Drosophila* Medium (Gibco, cat # 21720) supplemented with 10% FBS (Gibco) and 100 U/ml penicillin/streptomycin (Gibco). Cells at a 1×10^6^/ml density were transfected with 19 μg DNA constructs using 2 M CaCl_2_ and 2X HEPES-buffered saline (50 mM HEPES, 1.5 mM Na_2_HPO_4_, 280 mM NaCl, pH 7.1). Protein expression was verified by Western blotting (data not shown).

#### Flow cytometry

Transfected HEK 293T cells were harvested in 1 mM EDTA in PBS. The surface fraction of Venus-linked ASTN1 variants was labeled with rabbit anti-GFP (1:5,000; Invitrogen) for 20 minutes at 4 ^o^C followed by Alexa-647 donkey anti-rabbit (1:5,000; Life Technologies) for 25 minutes at 4 ^o^C. After each antibody incubation, the cells were washed in cold 10 % NDS in PBS. Cells were stained with Propidium Iodide (100 ng/ml; Sigma-Aldrich) for dead cell exclusion. Flow cytometry analysis on BD Accuri C6 (BD Biosciences) was carried out using 488 nm and 640 nm lasers and the CFlow Sampler software (BD Biosciences). A total of 20,000 single viable cells, identified by size and lack of Propidium Iodide staining, were analyzed per condition. Gates were set using non-transfected control cells and cells expressing cytosolic Venus, which were processed for live GFP labeling as described above. Data were analyzed by FlowJo v.9.3.3 (TreeStar Inc., Ashland, OR). Identical gates were applied to each condition.

#### Immunoprecipitation

Transfected HEK 293T cells or whole cerebella were extracted in ice-cold lysis buffer [50 mM Tris, pH 7.4, 150 mM sodium chloride, 0.5 % sodium deoxycholate, 1 % NP-40,1 mM EDTA and 1X protease inhibitor cocktail (Sigma-Aldrich)]. The extracts were triturated several times, incubated 20 minutes on ice and centrifuged at 14,000 rpm for 20 minutes at 4°C. The HEK 293T protein lysates (300–400 μg) or cerebellar protein lysates (700–750 μg) were then precleared with 25 μl Protein G/A Agarose beads (Calbiochem). After removing the beads, the lysates were incubated with 3 μg of a rabbit GFP antibody (Invitrogen), rabbit Astn1 antibody, or normal rabbit IgG (Santa Cruz Biotechnology) for 2 h at 4 ^o^C. Immunoprecipitates were collected on 50 μl Protein G/A Agarose beads by overnight rotation at 4 ^o^C, washed with lysis buffer and resuspended in 50 μl 2X Laemmli buffer. Western blotting was performed as described below.

#### Western blotting

Whole cerebella or purified cerebellar granule cells and Bergmann glial cells were extracted in ice-cold lysis buffer [50 mM Tris, pH 7.4, 150 mM sodium chloride, 0.5 % sodium deoxycholate, 1 % NP-40, 1 mM EDTA and 1X protease inhibitor cocktail (Sigma-Aldrich)]. The extracts were triturated several times, incubated 20 minutes on ice and centrifuged at 14,000 rpm for 20 minutes at 4°C. Protein concentrations were determined using the Pierce BCA Protein Assay Kit (Thermo Scientific). Protein samples in 2X Laemmli buffer were subjected to SDS-PAGE followed by transfer to Immobilon-P PVDF transfer membranes (Millipore). The membranes were blocked with 5% non-fat milk and labeled with primary and secondary antibodies in tris-buffered saline/0.1 % Tween, pH 7.6 (TBST), with each incubation for 1 h followed by washes in TBST. Primary antibodies were rabbit anti-GFP (1:10,000; Invitrogen) mouse anti-c-Myc (1:50; Calbiochem), mouse anti-N-cadherin (1:3000; BD Transduction Laboratories), rabbit anti-Astn1 (1:500) and mouse anti-GAPDH (1:10,000; Millipore). Secondary antibodies were horseradish peroxidase-conjugated anti-mouse (1:10,000; Jackson ImmunoResearch) or anti-rabbit (1:5,000; Jackson ImmunoResearch). Blots were developed with an ECL Western Blotting Detection kit (GE Healthcare) and exposed to Biomax XAR films (Carestream Health).

#### S2 cell adhesion assay

After transfecting the S2 cells for 24 h, 1.5×10^6^ cells from each transfection were mixed together in four conditions (Fig. 2) at a density of 3×10^6^ cells/well (1×10^6^ cells/ml) in ultra-low attachment 6-well plates (Corning Inc.). The cells were shaken gently at 28 ^o^C for up to 2 h to allow aggregation. Fab fragments of the Astn1 antibody were prepared and purified using Pierce Fab Micro Preparation Kit (Thermo Scientific) according to the manufacturer’s instructions. The Fab fragments were concentrated to 22 μg/ml using Amicon Ultra 2 ml Ultracel-10K centrifugal filters (Millipore) and were added to the S2 cells at a final concentration of 0.4 μg/ml (12). Cells were imaged on a Carl Zeiss Axiovert 135 fluorescent microscope with a 20X objective immediately after the conditions were set up (T = 0) and after 30 minutes, 1 h and 2 h. Longer incubations resulted in more dense cell cultures and aggregates forming in non-transfected control cells. Co-expression of the fluorophores, rather than tagged expression, ensured that they did not interfere with a potential *trans* binding. Cells expressing GFP (CDH2) or mCherry (control or ASTN1) were quantified in each aggregate and the proportion of mCherry-positive cells per aggregate was calculated for each condition and time point. The experiment was repeated six times.

#### Statistical analyses

The migration distance in the organotypic cultures (Fig. 5) was calculated by measuring the distance from the parallel fibers at the edge of the slice to the granule cell soma center using the “ruler” tool in Adobe Photoshop CS6. The number of migrating cells and non-migrating cells (round or multipolar) were quantified based on cell morphology, using the “count” tool in Adobe Photoshop CS6. In total, 700–1200 cells were quantified per control condition and 800–1300 cells per mutant condition from 5–7 slices per cerebellum (*n* = 3 per genotype). The neuron-glia distance (Fig. 6) was quantified in Adobe Photoshop CS6 by measuring the distance from the granule cell soma center to the nearest glial fiber. 30–45 cells were quantified for each condition and the experiment was repeated three times. Cerebellar size (Suppl Fig. S2) was measured using the freehand selection tool in ImageJ (National Institutes of Health) to encircle the whole cerebellum in midsagittal Nissl-stained sections of control and *Cdh2* cKO mice (*n* = 7 per genotype). For BrdU experiments (Fig. S3), the number of BrdU-positive cells in the EGL, ML, and IGL of the cerebellar cortex of control and *Cdh2* cKO mice (*n* = 4 per genotype) were counted using the “count” tool in Adobe Photoshop CS6 and the percentage in each layer determined. A total of 1600–2000 cells were counted per control mouse and 1500–1800 cells were counted per mutant mouse. Proliferating and apoptotic cells were quantified in phospho-histone H3 and Caspase-3 labeled sections of control and *Cdh2* cKO mice (*n* = 7 per genotype) by counting the cells in Adobe Photoshop CS6 and dividing the number of the cells with the length (proliferation) or with the area (apoptosis) of the cerebellar lobe. Microsoft Excel 2011 and IBM SPSS Statistics v21.0 were used for the data quantifications and statistical analyses. Differences between conditions were determined using unpaired *t*-tests for equal or unequal variances, except for migration distance in the slice cultures where Kruskal-Wallis and Mann-Whitney U non-parametric tests were used. Significance was set at *P* < 0.05 (two-sided). In the bar diagrams, data are presented as means with error bars representing the standard deviations. The migration distance data from the slice cultures are presented in box plots.

## Supplementary Figures

**Fig. S1.**
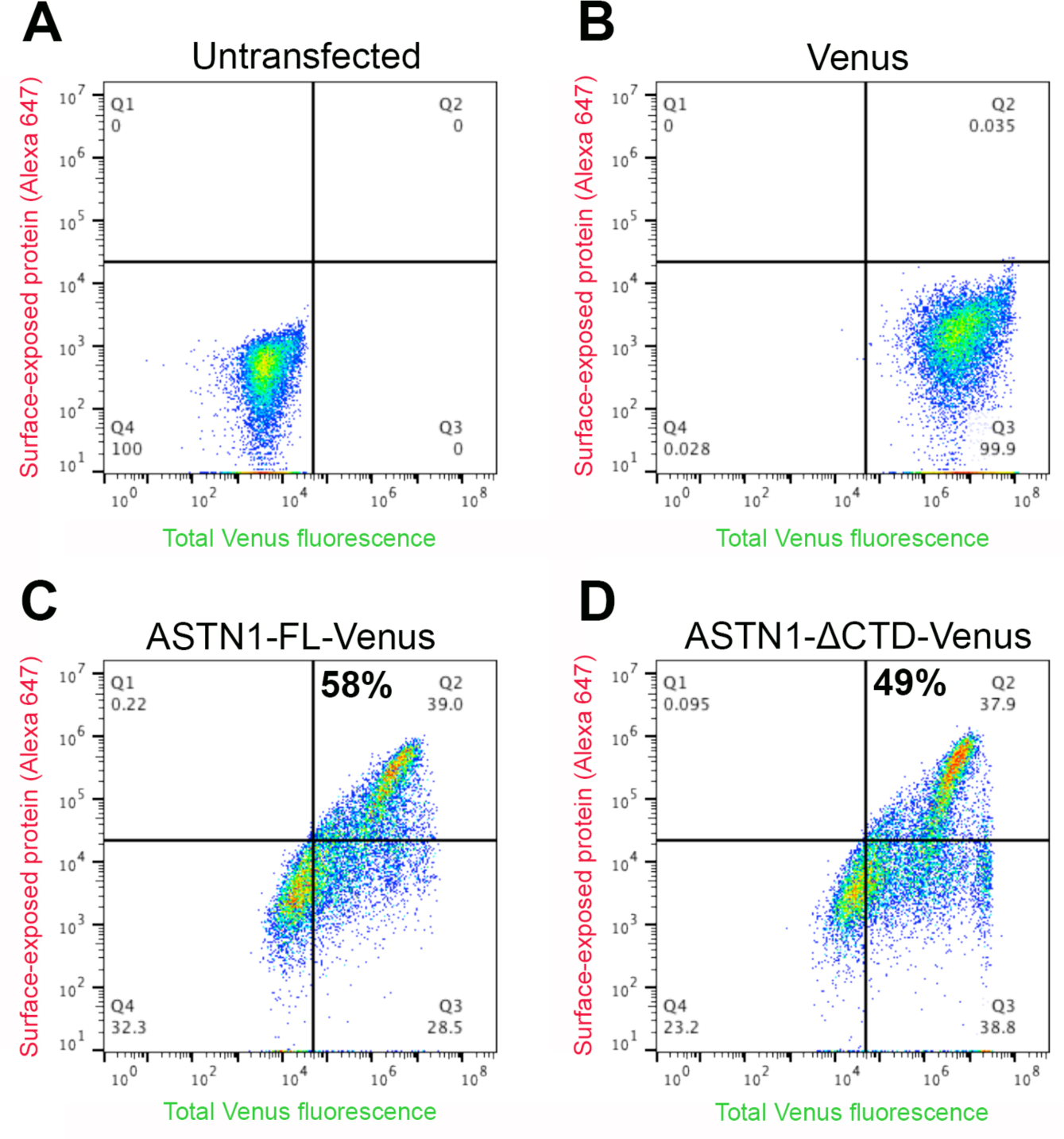
ASTN1-FL and ASTN1-ΔCTD are both expressed on the cell surface. Flow cytometry of live HEK 293T cells labeled with GFP and Alexa Fluor 647 antibodies. Control, untransfected cells **(A)** or cells transfected with *Venus* **(B)**, *Astnl-FL-Venus* **(C)**, or *Astnl-ΔCTD-Venus*, lacking the MACPF, FNIII and ANX-like domains in the C-terminus **(D)**. The x-axis shows total GFP fluorescence and the y-axis shows surface labeling (Alexa Fluor 647). Thus, cells expressing cytosolic Venus-tagged proteins are indicated in the lower right quadrant (Q3), while double positive cells (GFP+/Alexa Fluor 647+) in the upper right quadrant (Q2) express Venus-tagged proteins exposed on the cell surface. The percentage of cells expressing ASTN1 on the cell surface is depicted. Both ASTN1-FL and ASTNi-ΔCTD were significantly expressed on the cell surface (58 % and 49 % of live transfected cells, respectively).

**Fig. S2.**
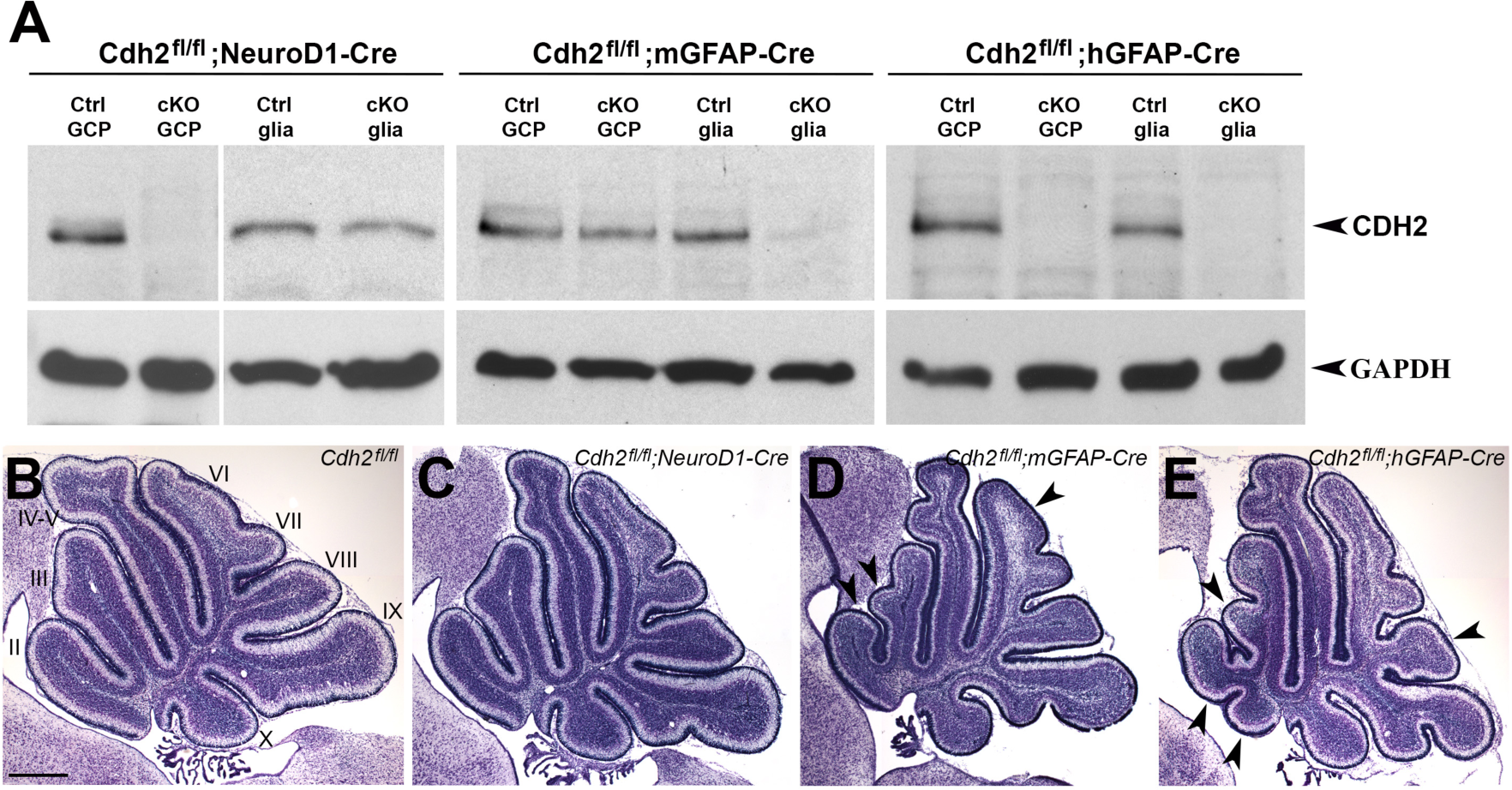
Conditional loss of CDH2 in cerebellar GCPs and/or BG. Western blots of protein lysates of GCPs and BG purified at P7 showing the absence of CDH2 expression in the three *Cdh2* ^*fl/fl*^ cKO lines. Loss of CDH2 was observed in GCPs from the *Cdh2* ^*fl/fl*^;*NeuroD1-Cre* mice, in BG from the *Cdh2* ^*fl/fl*^;*mGFAP-Cre* mice, and in both GCPs and glia from the *Cdh2* ^*fl/fl*^ *hGFAP-Cre* mice. Protein expression was compared to GAPDH. **(B - E)** Nissl staining of P7 cerebella of a *Cdh2*^*fl/fl*^ control mouse **(B)**, *Cdh2*^*fl/fl*^*;NeuroD1-Cre* mouse **(C)**, *Cdh2*^*fl/fl*^*;mGFAP-Cre* mouse (D) and *Cdh2*^*fl/fl*^; *hGFAP-Cre* mouse **(E)**. The cerebellar lobes are labeled with roman numerals in (B) Note the foliation defects in mice lacking *Cdh2* in BG or in both GCPs and BG (arrowheads in D, E). In particular, the ventral (I-III) lobes contained additional fissures resulting in extra lobules, while medio-dorsal (VI-VIII) lobes had fewer fissures and irregularly shaped folia. Scale bar represents 500 μm in (B - E).

**Fig. S3.**
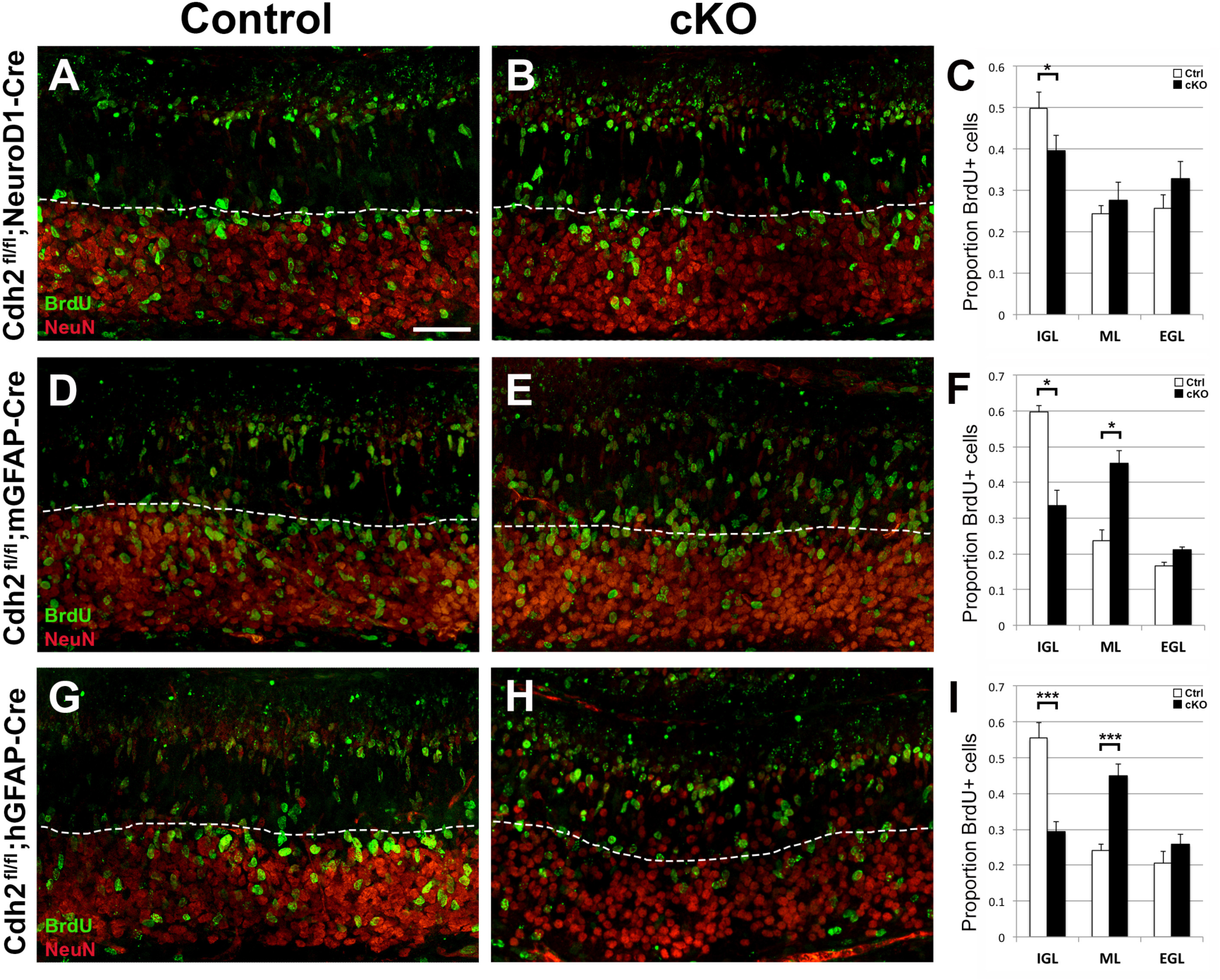
Birth-dating of GCs in *Cdh2* cKO mice. BrdU (green) and NeuN (red) labeling of P7 cerebella after BrdU injection at P5 (48 h). **(A - C)** Loss of *Cdh2* in GCPs in *Cdh2* ^*fl/fl*^;*NeuroD1-Cre* mice resulted in a 10 % decrease in the proportion of GCs reaching the IGL (dotted lines). **(D - F)** Severe migration defects were observed in *Cdh2* ^*fl/fl*^; *mGFAP-Cre* mice lacking *Cdh2* in BG, with a 27 % decrease in the proportion of GCs in the IGL and a 22 % increase in GCs in the ML. **(G - I)** Similar GCP migration defects were observed in the *Cdh2* ^*fl/fl*^;*hGFAP-Cre* mice, with a 26 % decrease in GCs in the IGL and a 21 % increase in GCs stalled in the ML. While the NeuN labeling showed a slightly more irregular IGL in the *Cdh2*^*fl/fl*^*;hGFAP-Cre* line, BrdU labeling revealed similar migration defects between the *Cdh2* ^*fl/fl*^;*mGFAP-Cre* and *Cdh2* ^*fl/fl*^; *hGFAP-Cre* lines, suggesting that the IGL patterning difference is dependent on the onset of the promoter (E13.5 for *hGFAP* and after P0 for *mGFAP)*. * *P* < 0.05; *** *P* < 0.001. Scale bar represents 50 μm in (A - H).

